# Pex14p phosphorylation modulates import of citrate synthase 2 into peroxisomes in *Saccharomyces cerevisiae*

**DOI:** 10.1101/2020.07.03.186833

**Authors:** Andreas Schummer, Renate Maier, Shiran Gabay-Maskit, Tobias Hansen, Wignand W. D. Mühlhäuser, Ida Suppanz, Amir Fadel, Maya Schuldiner, Wolfgang Girzalsky, Silke Oeljeklaus, Einat Zalckvar, Ralf Erdmann, Bettina Warscheid

## Abstract

The peroxisomal biogenesis factor Pex14p is an essential component of the peroxisomal matrix protein import machinery. Together with Pex13p and Pex17p, it is part of the membrane-associated peroxisomal docking complex in yeast, facilitating the binding of cargo-loaded receptor proteins for translocation of cargo proteins into the peroxisome. Furthermore, Pex14p is part of peroxisomal import pores. The central role of Pex14p in peroxisomal matrix protein import processes renders it an obvious target for regulatory mechanisms such as protein phosphorylation. To explore this possibility, we examined the state of Pex14p phosphorylation in *Saccharomyces cerevisiae*. Phos-tag-SDS-PAGE of Pex14p affinity-purified from solubilized membranes revealed Pex14p as multi-phosphorylated protein. Using mass spectrometry, we identified 16 phosphorylation sites, with phosphorylation hot spots located in the N- and C-terminal regions of Pex14p. Analysis of phosphomimicking and nonphosphorylatable variants of Pex14p revealed a decreased import of GFP carrying a peroxisomal targeting signal type 1, indicating a functional relevance of Pex14p phosphorylation in peroxisomal matrix protein import. We show that this effect can be ascribed to the phosphomimicking mutation at serine 266 of Pex14p (Pex14p-S266D). We further screened the subcellular distribution of 23 native GFP-tagged peroxisomal matrix proteins by high-content fluorescence microscopy. Only Cit2p, the peroxisomal isoform of citrate synthase, was affected in the Pex14p-S266D mutant, showing increased cytosolic localization. Cit2p is part of the glyoxylate cycle, which is required for the production of essential carbohydrates when yeast is grown on non-fermentable carbon sources. Pex14p-S266 phosphosite mutants showed reversed growth phenotypes on oleic acid and ethanol with acetyl-CoA formed in peroxisomes and the cytosol, respectively. Our data point to the control of the peroxisomal import of Cit2p via the state of Pex14p phosphorylation at S266, which may help *S. cerevisiae* cells to rapidly adjust their carbohydrate metabolism according to the nutritional conditions.

## Introduction

Peroxisomes are highly dynamic metabolic organelles that are present in nearly all eukaryotic cells and fulfill a wide variety of metabolic functions, depending on organism, tissue, and environmental condition. Common metabolic functions include β-oxidation of fatty acids, which in the yeast *Saccharomyces cerevisiae* exclusively occurs in peroxisomes (Kunau et al., 1988; Poirier et al., 2006), and the degradation of hydrogen peroxide, generated in peroxisomes during various oxidative reactions (Wanders and Waterham, 2006b). In addition, peroxisomes are involved in a series of species- or tissue-specific metabolic activities such as the glyoxylate cycle in yeast and plants (Breidenbach and Beevers, 1967; Kunze et al., 2006), glycolysis in Trypanosoma (Opperdoes and Borst, 1977), the synthesis of plasmalogens and bile acids in mammals (Wanders and Waterham, 2006a), or the generation of penicillin in filamentous fungi (Müller et al., 1991). Maintenance of the highly dynamic peroxisomal system and its adjustment to varying physiological conditions rely on several processes including peroxisomal protein targeting, import of matrix proteins and peroxisomal membrane biogenesis proteins as well as growth and division, and turnover of peroxisomes by autophagy. All these processes need to be well coordinated and tightly regulated to ensure the functionality of the peroxisomal system.

Since peroxisomes do not contain their own DNA, all peroxisomal matrix proteins are synthesized in the cytosol and need to be imported into the organelle (Walter and Erdmann, 2019). For their correct sorting to the peroxisomal matrix, most proteins possess a peroxisomal targeting signal (PTS), either a PTS1 located at the C-terminus or a PTS2 within the N-terminus (Gould et al., 1989; Swinkels et al., 1991; Brocard and Hartig, 2006). PTS1 and PTS2 are recognized in the cytosol by their cognate receptors Pex5p or Pex9p and Pex7p, respectively, with the latter in *S. cerevisiae* additionally requiring the co-receptors Pex18p or Pex21p (Effelsberg et al., 2016; Yifrach et al., 2016; Walter and Erdmann, 2019). At the peroxisomal membrane, cargo-loaded receptors bind to a docking complex consisting of Pex14p and Pex13p, completed by Pex17p in yeast (Elgersma et al., 1996; Erdmann and Blobel, 1996; Gould et al., 1996; Albertini et al., 1997; Huhse et al., 1998; Azevedo and Schliebs, 2006). Upon docking, the cargo protein is translocated across the peroxisomal membrane into the matrix. At the end of the import cycle, the receptor is extracted from the peroxisomal membrane in sequential ATP- and ubiquitin-dependent steps (Platta et al., 2014). In *S. cerevisiae*, two distinct pores for the PTS1 and PTS2 import pathways have been identified, both comprising Pex14p as an essential pore-forming element together with either the PTS1 receptor Pex5p or the PTS2 co-receptor Pex18p and Pex17p (Meinecke et al., 2010; Montilla-Martinez et al., 2015).

Considering the prominent role of Pex14p in peroxisomal import pathways, it represents an obvious point of regulation for peroxisomal matrix protein import. In fact, Pex14p of *S. cerevisiae* has been reported to be phosphorylated in several global phosphoproteomic studies (summarized in Oeljeklaus et al., 2016; Albuquerque et al., 2008; Gnad et al., 2009; Holt et al., 2009; Yachie et al., 2011; Hu et al., 2019). However, information about the biological relevance of Pex14p phosphorylation is lacking.

In this work, we established a comprehensive Pex14p *in vivo* phosphorylation map using high-resolution mass spectrometry (MS) and report 16 high-confidence phosphorylation sites. We generated phosphomimicking and non-phosphorylatable Pex14p mutants, in which single serine or threonine residues were substituted with alanine or aspartate, and systematically analyzed all phosphosite mutants by fluorescence microscopy. Our data show that the peroxisomal import for GFP carrying a PTS1 is reduced in cells that express the phosphomimicking variant of serine 266, which is located within in the receptor-binding region of Pex14p. In addition, these cells are impaired in growth on oleic acid, for which peroxisomes are essential. A screen of 23 GFP-tagged native peroxisomal matrix proteins revealed that cells expressing Pex14p-S266D exhibit a reduced peroxisomal import for the peroxisomal isoform of citrate synthase (Cit2p), a protein involved in the glyoxylate cycle, which we confirmed by biochemical analysis. Furthermore, the S266D mutant shows wild-type-like growth on ethanol when acetyl-CoA is formed in the cytosol, but growth of the S266A mutant is impaired. Our data suggest that site-specific phosphorylation of Pex14p at S266 provides a means to rapidly modulate the peroxisomal import and thus the subcellular localization of Cit2p in response to the metabolic state of the cell.

## Material and methods

### Strains, media, and growth conditions

*S. cerevisiae* strains used in this study as well as primers and DNA templates used for genomic manipulation of target strains are listed in **Supplementary Table S1**. Sequences of primers used for the amplification of integration cassettes are specified in **Supplementary Table S2**. The strain SC38 was generated by gene disruption using pUG27 or pUG73 followed by marker rescue using pSH47 (containing Cre recombinase for LoxP recombination) as described before (Güldener et al., 1996). Deletions of *PEX14* were introduced according to Goldstein et al. (1999). Yeast strains expressing Pex14p genomically tagged at its C-terminus with the tobacco etch virus protease cleavage site (TEVcs) and Protein A (Pex14p^TPA^) as well as genomically integrated TPA-tagged Pex14p phosphosite mutants were generated by transforming haploid yeast cells (Schiestl and Gietz, 1989; Becker and Guarente, 1991) with PCR products obtained as described before (Knop et al., 1999). Transformants were selected for the appropriate marker for negative selection or, if applicable, by adding 5-fluoroorotic acid to the agar for positive selection (Boeke et al., 1984). All chromosomal manipulations were confirmed by PCR. Chromosomal tagging and genomic integration of phosphosite mutants were further verified by sequencing of PCR products and/or immunoblotting.

Minimal liquid medium contained 0.17% (w/v) yeast nitrogen base (YNB) without amino acids, 0.5% (w/v) ammonium sulfate, 0.3% (w/v) glucose, and selected amino acids and nucleobases (pH 6.0) as described before (Oeljeklaus et al., 2012). YNO medium consisted of 0.1% (w/v) oleic acid, 0.05% (v/v) Tween 40, 0.17% (w/v) YNB without amino acids, 0.5% (w/v) ammonium sulfate, and selected amino acids and nucleobases (pH 6.0).

To induce peroxisome proliferation, cells were grown at 30°C in minimal medium until they reached an optical density at 600 nm (OD_600_) of 1 - 1.5, shifted to YNO medium and cultivated for further 16 hours or as indicated. For growth of cells in glucose or ethanol medium, minimal medium was supplemented with 2% (w/v) glucose or 2% (v/v) ethanol.

### Plasmids and cloning techniques

Plasmids, primers, restriction enzymes, DNA templates and target plasmids used for cloning are listed in **Supplementary Table S3**; sequences of the primers are specified in **Supplementary Table S2**. Plasmids were amplified in *Escherichia coli* DH5α or TOP10 following standard protocols and generated using standard cloning techniques (Sambrook et al., 1989). Individual serine or threonine residues in Pex14p were exchanged to alanine or aspartate by site-directed mutagenesis (Papworth et al., 1996). To simultaneously insert multiple mutations, customized DNA was obtained (Thermo Fisher Scientific/GeneArt; Eurofins) and cloned into *PEX14* using restriction enzymes (see **Supplementary Table S3** for more details).

### Two-hybrid analyses

The yeast reporter strain PCY2 *pex14Δ* was transformed with two-hybrid plasmids pPC86 and pPC97 (Chevray and Nathans, 1992) or derivatives thereof and grown on synthetic medium lacking tryptophan and leucine for 3 days at 30°C. ß-galactosidase activity of transformed cells was determined by a filter assay as described before (Rehling et al., 1996) using X-Gal as substrate. For quantification, β-galactosidase activity was determined by performing a liquid assay using 2-nitrophenyl β-D-galactopyranoside as substrate according to manufacturer’s instructions (Clonetech, Yeast Protocols Handbook, 2009).

### Affinity purification of Pex14p^TPA^

Pex14p was affinity-purified from crude membrane fractions prepared from oleic acid-induced Pex14p^TPA^-expressing cells as described before with minor modifications (Agne et al., 2003; Oeljeklaus et al., 2012). Cells were harvested by centrifugation, washed with deionized water, and resuspended in lysis buffer supplemented with protease and phosphatase inhibitors (20 mM Tris, 80 mM NaCl, pH 7.5, 174 μg/ml PMSF, 2 μg/ml aprotinin, 0.35 μg/ml bestatin, 1 μg/ml pepstatin, 2.5 μg/ml leupeptin, 160 μg/ml benzamidine, 5 μg/ml antipain, 6 μg/ml chymostatin, 420 μg/ml NaF). Lysates were prepared using glass beads (Lamb et al., 1994), cell debris and beads were removed, and the lysate was centrifuged for 90 min at 100,000 x g and 4°C. The pellet was resuspended in lysis buffer containing 10% (v/v) glycerol. For phosphosite analysis, the protein concentration was adjusted to 5 mg/ml and proteins were solubilized using 1% (v/v) Triton X-100. For the analysis of native Pex14p complexes, the protein concentration was adjusted to 3.35 mg/ml and solubilization was performed with 1% (v/v) digitonin. The detergent extract was cleared by centrifugation (100,000 x g, 1 h, 4°C) and the supernatant was used to affinity-purify Pex14p^TPA^ via IgG Sepharose. Proteins were eluted by incubation with 0.2 M glycine (pH 2.4) or TEV protease. The TEV protease, carrying a His tag, was removed from the eluate by adding Ni-NTA agarose. Eluted proteins were collected by centrifugation and either used immediately (for dephosphorylation) or precipitated by adding four volumes of ice-cold acetone followed by incubation at −20°C for at least 1 h (for immunoblot and MS analyses).

### Dephosphorylation of Pex14p and Phos-tag SDS-PAGE

Pex14p^TPA^ affinity-purified from Triton X-100 extracts was mixed with 10x phosphatase buffer (0.5 M HEPES, 1 M NaCl, 20 mM DTT, 0.1% Brij 35, pH 7.5), MnCl_2_ (1 mM final concentration), and 800 units of lambda protein phosphatase (λ-PPase; New England Biolabs) and incubated for 20 min at 30°C under slight agitation. As control, the reaction was performed without λ-PPase using the same amount of Pex14p^TPA^. Reactions were stopped by adding SDS sample buffer and proteins were separated by Phos-tag SDS-PAGE using a standard 10% SDS polyacrylamide gel additionally containing 50 µM Phos-tag reagent (Phos-tag acrylamide AAL-107; Wako Pure Chemical Industries, Ltd., Japan) and 100 mM MnCl_2_ in the separation gel. Electrophoresis was carried out at 80 V for 4 h. Subsequently, the gel was incubated for 20 min in Western blot transfer buffer (25 mM Tris, 192 mM glycine, 20% [v/v] methanol) containing 10 mM EDTA to quench the Phos-tag reagent and washed for 10 min in transfer buffer without EDTA. Proteins were transferred to a polyvinylidene difluoride membrane at a constant voltage of 30 V and 4°C for 14 h.

### Sedimentation assay

Spheroplasting of yeast cells was performed as described before (Erdmann and Blobel, 1995) with slight modifications. The cell wall was digested by adding lyticase at a final concentration of 1,000 units per g of cells (wet weight). Cells were homogenized, and unbroken cells, cell debris and nuclei were removed by centrifugation (2x 10 min, 600 x g, 4°C). The resulting postnuclear supernatant (PNS) was separated into an organellar pellet and the cytosolic fraction by centrifugation (25,000 x g, 20 min, 4°C) through a 0.5-M sucrose cushion.

### Whole yeast cell lysates

Cells were harvested, washed with deionized water, and lysed by TCA precipitation as described previously (Platta et al., 2004). Following centrifugation (10 min, 21,000 x g), precipitates were resuspended in 1% (w/v) SDS/0.1 M NaOH and analyzed by SDS-PAGE according to standard protocols.

### Immunoblotting and antibodies

Immunoblot analyses were performed according to Harlow and Lane (Harlow and Lane, 1988) with polyclonal rabbit antibodies raised against Pcs60p (Blobel and Erdmann, 1996), Pex5p (Albertini et al., 1997), Pex7p (Stein et al., 2002), Pex13p (Girzalsky et al., 1999), Pex14p (Albertini et al., 1997), Pex17p (Huhse et al., 1998), and Protein A (Sigma-Aldrich, cat. # P3775). In addition, the goat polyclonal antibody anti-Cta1p (Kragler et al., 1993) and the mouse monoclonal antibodies anti-GFP (Roche Diagnostics, cat. # 11814460001), anti-Pgk1p (Thermo Fisher Scientific, cat. # 459250), anti-Por1p (Thermo Fisher Scientific, cat. # 459500) and anti-Protein A (Sigma-Aldrich, cat. # P2921) were used. Primary antibodies were detected with horseradish peroxidase-conjugated anti-rabbit, anti-goat and anti-mouse antibodies (Sigma-Aldrich, cat. # A0545, A8919, and A9044, respectively). Chemiluminescence signals were detected using the ECL2 Western Blotting Substrate or SuperSignal West Femto Maximum Sensitivity Substrate (Thermo Fisher Scientific, Waltham, USA) and a ChemoCam Camera System (INTAS Science Imaging Instruments GmbH, Göttingen, Germany). Image processing was restricted to cropping, scaling and contrast adjustment using Adobe Photoshop CS5 (v. 12.0.4 x64). Immunoblot signals were quantified using the software Quantity One (version 4.6.9.; Bio-Rad, Hercules, USA).

### Sample preparation for fluorescence microscopy

Yeast cells were grown for 16 hours in glucose- or oleic acid-containing medium. Aliquots of 5.4 ml per culture were taken and cells were fixed by directly adding formaldehyde to a final concentration of 3.7% (v/v). Samples were incubated for 10 min at RT with slight agitation. Cells were harvested by centrifugation (5 min at 2,000 x g), resuspended in 1 ml of potassium phosphate buffer (0.1 M, pH 6.8) containing 3.7% (v/v) formaldehyde and incubated for 1 h at RT with slight agitation. Cells were harvested again (30 s at 15,700 x g) and resuspended in 1 ml of potassium phosphate buffer (0.1 M, pH 6.8) containing 10 mM ethanolamine. Following incubation for further 10 min at RT, cells were washed twice with phosphate-buffered saline (PBS) and resuspended in 100 - 200 µl of PBS containing 0.1% (v/v) Triton X-100.

### Fluorescence microscopy

Wide-field fluorescence microscopy was performed using a Zeiss Axio Observer.Z1 microscope (Zeiss) equipped with an alpha Plan-Aprochromat 100x/1.45 oil objective and an AxioCam HRm Rev.3 camera. GFP signals were visualized using a 38 HE filterset from Zeiss. Z-stacks containing approx. 20 images were acquired using an exposure time of 600 ms at a depth of focus of 0.73 µm. Images were acquired using the Zen software (version 10.0.14393; Zeiss) and processed using Adobe Photoshop CS5 (version 12.0.4 x64; Adobe Systems Incorporated) applying the same parameters to all images. Images in **Figures 2B, 3A, 5B**, and **S5** show a single focal plane. Image analysis was performed in independent replicates and data were assessed manually by two different researchers in blind experiments to exclude bias in image analysis. To show cell boundaries, bright field images were adjusted such that only the cell boundaries were visible and subsequently added into the blue channel using Photoshop (Motley and Hettema, 2007).

For the quantification of the cytosolic fluorescence of the images shown in **Figures 5D** and **S5B** (i.e. Pex14p wild-type, S266A and S266D cells), a slice of the z-series was selected from the bright-field channel of the images. Selection was based on best visual separation of individual cells. For cell segmentation, the Python package skimage was used. In short, binary thresholding of bright field images followed by removal of small objects (< 300 pixels), closing of small openings (disk size of 5 pixels), and filling of open holes were sufficient for cell segmentation. Cartesian coordinates of a rectangular area encompassing the cell were extracted. Of the GFP z-series, a maximum intensity z-projection was performed in Fiji/ImageJ (version 1.52p; Schindelin et al., 2012). Afterwards, Cartesian coordinates of the bright field channel of segmented cells were mapped onto the respective GFP intensity projection. To remove false positives from the segmented cells, images with an axis of > 100 pixels or < 40 pixels, reaching the maximum intensity of 255 (grayscale image), or having a standard deviation of intensities of < 5 were removed. The lower quantile of single cell intensities was used as surrogate for the cytosolic GFP intensity and the mean value of the lowest 10% intensities as background. Images in which the cytosolic GFP was located too close to a border of an image (5%) were considered incomplete and also removed prior to quantitative analysis. The cytosolic intensities were corrected for the background and median-centered for each strain. To test for statistically significant differences in the cytosolic fluorescence of Pex14p wild-type, S266A and S266D cells, a Welsh’s t-test was performed with n = 73 for wild-type, n = 69 for S266A, and n = 174 for S266D for GFP-Cit2p cells (**Figure 5E**) and n = 33 for wild-type, n = 48 for S266A, and n = 20 for S266D for GFP-Mdh3p cells (**Figure S5C**).

### Yeast library preparation

Query strains were constructed on the basis of an automated mating compatible strain. The query strains contained a peroxisomal marker (Pex3p-mCherry::HIS) and a *PEX14*-TEVcs-ProteinA::KanMX4 cassette integrated into the *PEX14* locus. The *PEX14* gene was either wild-type or mutated in S266A or S266D. Using an automated mating method (Tong and Boone, 2006; Cohen and Schuldiner, 2011), the query strains were crossed with 23 strains that in each of them one protein (Pex5p cargo proteins, PTS2 proteins and Pnc1p, which piggybacks on a PTS2 protein) was expressed under a generic constitutive *SpNOP1* promoter and N-terminally tagged with GFP (collected from the *SpNOP1* GFP-peroxi library [Dahan et al., 2017]). To perform the manipulations in high-density format, we used a RoToR bench top colony arrayer (Singer Instruments). In short: mating was performed on rich medium plates, and selection for diploid cells was performed on SD(MSG)-HIS-URA plates containing kanamycin (200 μg/ml). Sporulation was induced by transferring cells to nitrogen starvation media plates for 7 days. Haploid cells containing the desired mutations were selected by transferring cells to SD-URA-HIS-LYS-ARG-LEU plates containing antibiotics at the same concentrations as above, alongside the toxic amino acid derivatives canavanine and thialysine (Sigma-Aldrich) to select against remaining diploids and select for spores with an α mating type.

### Automated high-throughput fluorescence microscopy

The yeast collections were visualized using an automated microscopy setup as described previously (Breker et al., 2013). In short: cells were transferred from agar plates into 384-well polystyrene plates (Greiner) for growth in liquid media using the RoToR arrayer robot. Liquid cultures were grown in a LiCONiC incubator, overnight at 30°C in SD-URA-HIS medium. A JANUS liquid handler (PerkinElmer) connected to the incubator was used to dilute the strains to an OD_600_ of ∼0.2 into plates containing SD medium (6.7 g/l yeast nitrogen base and 2% glucose) supplemented with complete amino acids. Plates were incubated at 30°C for 4 h. The cultures in the plates were then transferred by the liquid handler into glass-bottom 384-well microscope plates (Matrical Bioscience) coated with concanavalin A (Sigma-Aldrich). After 20 min, cells were washed three times with SD-riboflavin complete medium to remove non-adherent cells and to obtain a cell monolayer. The plates were then transferred to the ScanR automated inverted fluorescent microscope system (Olympus) using a robotic swap arm (Hamilton). Images of cells in the 384-well plates were recorded in the same liquid as the washing step using a 60x air lens (NA 0.9) and with an ORCA-ER charge-coupled device camera (Hamamatsu). Images were acquired in two channels: GFP (excitation filter 490/20 nm, emission filter 535/50 nm) and mCherry (excitation filter 572/ 35 nm, emission filter 632/60 nm). All images were taken at a single focal plane using the same exposure time. Raw images were subsequently processed and analyzed using Fiji/ImageJ (version 1.51q; Schindelin et al., 2012) applying the same parameters to ensure that any differences observed in distribution are not a result of differential processing of images.

### Mass spectrometric analyses

For *in vivo* phosphosite analyses, affinity-purified and acetone-precipitated Pex14p was proteolytically digested with trypsin (Promega), AspN (Sigma-Aldrich), or Lys-C (Promega). Proteins were resuspended in 100 µl of the appropriate digestion buffer (trypsin: 60% [v/v] methanol/20 mM NH_4_HCO_3_; AspN: 100 mM NH_4_HCO_3_; Lys-C: 25 mM Tris-HCl, pH 8.5, containing 1 mM EDTA) and incubated overnight at 37°C with 225 ng trypsin, 160 ng AspN, or 120 ng Lys-C. For the analysis of native Pex14p complexes affinity-purified from digitonin-solubilized membrane fractions, proteins were separated by SDS-PAGE on a 4-12% NuPAGE BisTris gradient gel (Thermo Fisher Scientific/Life Technologies) and stained using colloidal Coomassie Brilliant Blue. Gel lanes were cut into 14 equal slices, cysteine residues were reduced (5 mM Tris[2-carboxy-ethyl]phosphine/10 mM NH_4_HCO_3_; 30 min at 37°C) and alkylated (50 mM chloroacetamide/10 mM NH_4_HCO_3_; 30 min at room temperature), and proteins were in-gel digested with trypsin (66 ng per slice, 37°C, overnight). Peptide mixtures were dried *in vacuo* and peptides were resuspended in 0.1% trifluoroacetic acid prior to MS analysis.

Liquid chromatography-mass spectrometry (LC-MS) analyses of Pex14p phosphosites were performed with an UltiMate 3000 RSLCnano HPLC system (Thermo Fisher Scientific, Dreieich, Germany) coupled to either an LTQ-FT or an LTQ-Orbitrap XL instrument (Thermo Fisher Scientific, Bremen, Germany). RSLC systems were equipped with C18 μ-precolumns (PepMap™, 5 mm x 0.3 mm; Thermo Fisher Scientific, Bremen, Germany) and C18 reversed-phase nano RSLC columns (Acclaim PepMap™, 25 cm x 75 μm, 2 μm particle size, 100 Å pore size; Thermo Fisher Scientific) and operated at a temperature of 40°C. MS instruments were equipped with a nanoelectrospray ion source and stainless steel emitters (Thermo Fisher Scientific; LTQ-FT) or distal coated Silica Tips (FS360-20-10-D, New Objective, Woburn, USA; LTQ-Orbitrap) and externally calibrated using standard compounds. Peptide fragmentation for Pex14p phosphosite analysis was generally performed on multiply charged precursor ions by multistage activation (MSA; Schroeder et al., 2004; Boersema et al., 2009) with neutral loss masses of 32.7, 49 and 98 Da and applying a normalized collision energy of 35%, an activation q of 0.25, and an activation time of 30 ms. The dynamic exclusion time for previously fragmented precursor ions was set to 45 s. Proteolytic Pex14p peptides analyzed at the LTQ-FT were separated using a binary solvent system consisting of 0.1% formic acid (FA) (solvent A) and 86% acetonitrile (ACN)/0.1% FA (solvent B) at a flow rate of 300 nl/min. Peptides were eluted with a gradient of 4 - 34% solvent B in 35 min followed by 34 - 82% B in 5 min and 5 min at 82% B. Full MS scans in the range of *m/z* 370 - 1,700 were acquired in the ion cyclotron-resonance cell at a resolution (R) of 25,000 (at *m/z* 400) with an automatic gain control (AGC) of 2 × 10^6^ ions and a maximum fill time of 500 ms. Up to 5 of the most intense multiply charged precursor ions (TOP5 method) were selected for MSA fragmentation in the linear ion trap applying the following parameters: signal threshold, 500; AGC, 3 × 10^4^ ions; max. fill time, 500 ms. Proteolytic peptides analyzed at the LTQ-Orbitrap XL were separated employing the solvent system and the flow rate described above. Peptides were eluted with the following gradient: 1 - 30% B in 75 min followed by 30 - 45% B in 30 min, 45 - 70% B in 25 min, 70 - 99% B in 5 min and 5 min at 99% B. Full MS scans (*m*/*z* 370 - 1,700) were acquired in the orbitrap applying the following parameters: R, 60,000 (at *m*/*z* 400); AGC, 2 × 10^6^ ions; max. fill time, 500 ms. A TOP6 method was applied for peptide fragmentation in the linear ion trap with the following parameters: signal threshold, 2,500; AGC, 10^4^ ions; max. fill time, 400 ms.

LC-MS analyses of native Pex14p complexes were performed using an UltiMate 3000 RSLCnano HPLC system/Orbitrap Elite (Thermo Fisher Scientific, Bremen, Germany) system. The LC system was equipped with nanoEase™M/Z Symmetry C18 trap columns (100 Å pore size, 5 µm particle size, 20 × 0.18 mm; Waters, Milford, USA) and a 25 cm x 75 µm nanoEase™M/Z HSS C18 T3 reversed-phase nano LC column (100 Å pore size, 2 µm particle size; Waters) and operated at 40°C. Peptides were eluted with a solvent system consisting of 0.1% FA/4% dimethyl sulfoxide (DMSO) (solvent A) and 48% methanol/30% ACN/0.1% FA/4% DMSO (solvent B). The gradient was as follows: 1 - 65% solvent B in 30 min, 65 - 80% B in 5 min and 3 min at 80% B at a flow rate of 300 nl/min. The MS instrument was equipped with a nanoelectrospray ion source and distal coated Silica Tips and was externally calibrated with standard compounds. Peptide fragmentation was performed on multiply charged precursor ions by collision-induced dissociation (CID) with a normalized collision energy of 35%, an activation q of 0.25, an activation time of 30 ms, and a dynamic exclusion of 45 s. Full MS scans in the range of *m*/*z* 370 - 1,700 were acquired in the orbitrap with the following parameters: R, 120,000 (at *m*/*z* 400); AGC, 10^6^ ions; max. fill time, 200 ms. A TOP25 method was employed for CID peptide fragmentation in the linear ion trap applying a signal threshold of 2,500, and AGC of 5,000 ions, and a max. fill time of 150 ms.

### Mass spectrometric data analysis

For processing of mass spectrometric raw data, the software MaxQuant (version 1.5.2.8; Cox and Mann, 2008) and its integrated search engine Andromeda (Cox et al., 2011) were used. Peaklists of MS/MS spectra, generated by MaxQuant using default settings, were searched against custom-made protein databases. For the datasets obtained to identify *in vivo* phosphorylation sites of affinity-purified wild-type Pex14p, a modified version of the *Saccharomyces* Genome Database (SGD; 6,750 entries; downloaded 02/03/2011) was used in which the entry for Pex14p was replaced with an extended Pex14p sequence containing at its C-terminus the residues of the TPA tag (RTLQVDGSENLYFQ) that remain after TEV cleavage. For the analysis of proteins in native Pex14p complexes (wild-type Pex14p, Pex14p-S266A or -S266D), the database additionally contained the sequences of the S266A and S266D variants. Common contaminants provided by Max Quant as well as the sequences of Protein A and the TEV protease were included in all searches. Database searches were performed with the appropriate enzymatic specificity (*i*.*e*. trypsin, AspN, or Lys-C) allowing four missed cleavages. The max. molecular mass was set to 6,000 Da, mass tolerances for precursor ions were 20 ppm for the first and 6 ppm for the main search and 0.5 Da for fragment ions. Oxidation of methionine, acetylation of protein N-termini and, for the identification of phosphosites, phosphorylation of serine, threonine and tyrosine residues were set as variable modifications. For peptide identification, a minimum length of six amino acids was required and an FDR of 1% was applied on both peptide and protein level. The options ‘match between runs’ and, for the quantitative analysis of proteins present in native Pex14p complexes, ‘iBAQ’ (abbreviation for ‘intensity-based absolute quantification’) were enabled.

To be reported as confident Pex14p *in vivo* phosphosites, phosphopeptides needed to be identified with a posterior error probability (PEP) of < 0.01 and the MaxQuant localization probability for individual phosphosites needed to be at least 0.95. For a complete list of all Pex14p peptides identified in these datasets (phosphorylated and non-phosphorylated; four independent replicates), see **Supplementary Table S4**. The correct assignment of phosphosites to a distinct residue was further validated by manual inspection of MS/MS spectra.

To assess and compare the abundance of proteins in native protein complexes of Pex14p^TPA^ wild-type and its S266A and S266D variants, iBAQ intensities of individual proteins were normalized to the summed iBAQ value of all proteins identified in each replicate, and the value for Pex14p was set to 100%. We determined mean values of two independent replicates (see **Supplementary Table S5** for a list of proteins identified and the respective iBAQ values).

## Results

### Pex14p is multiply phosphorylated *in vivo*

With the aim to obtain further insight into the regulation of peroxisomal matrix protein import, we studied phosphorylation of Pex14p in *S. cerevisiae*. To establish a most comprehensive Pex14p *in vivo* phosphorylation map, we affinity-purified TPA-tagged Pex14p expressed from its native chromosomal location from a crude membrane fraction of yeast cells grown under peroxisome-proliferating conditions (Veenhuis et al., 1987). The obtained eluate fraction was analyzed by phosphate-affinity (Phos-tag) SDS-PAGE, in which phosphorylated proteins migrate slower than their non-phosphorylated counterparts (Kinoshita et al., 2006). Immunoblot analysis revealed the occurrence of different Pex14p phospho-isoforms in the higher molecular weight region, which indicates that native Pex14p is phosphorylated at multiple amino acid residues (**Figure 1A**). Substantiating this conclusion, treatment of Pex14p with λ-phosphatase resulted in a single Pex14p signal and a shift in electrophoretic mobility reflecting faster migration of dephosphorylated Pex14p.

**Figure 1.**
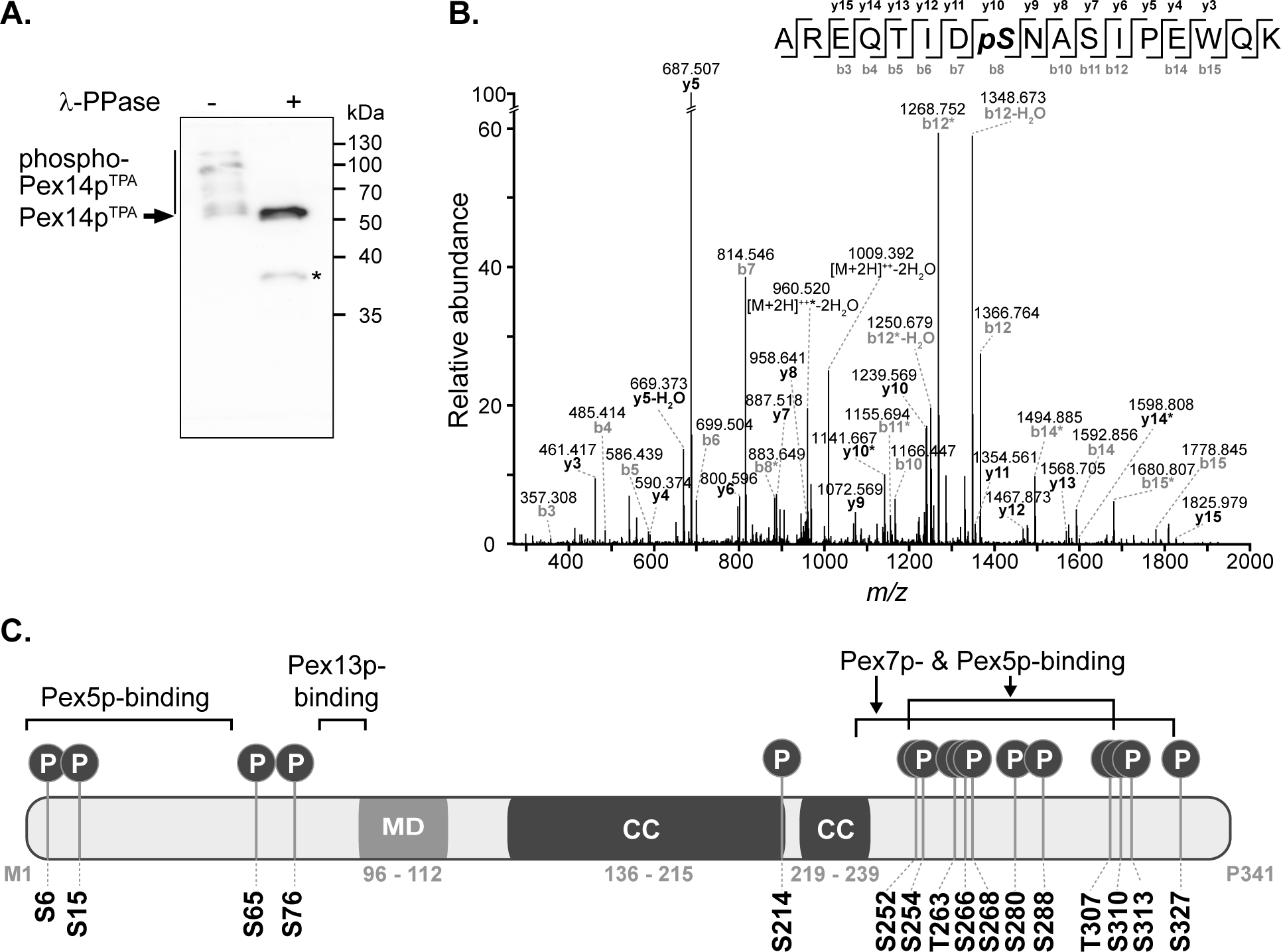
Pex14p is multiply phosphorylated *in vivo*. (**A**) TPA-tagged Pex14p (Pex14p^TPA^) expressed from its native chromosomal location was affinity-purified from a Triton X-100-solubilized crude membrane fraction of yeast cells grown in oleic acid-containing medium. Affinity chromatography was performed using IgG-coupled Sepharose and proteins were eluted with glycine (pH 2.4). Proteins from phosphatase (λ-PPase)-treated (+) and untreated (-) samples were analyzed by Phos-tag SDS-PAGE and immunoblotting using anti-Pex14p antibodies. Note that also non-phosphorylated proteins generally migrate slower in Phos-tag gels compared to normal SDS gels (Kinoshita et al., 2009). *, Pex14p degradation band. (**B**) Representative MS/MS spectrum showing phosphorylation of Pex14p at S310 (AREQTIDpSNASIPEWQK; *m/z* 1027.474; z = +2). *, fragment ion with neutral loss of H_3_PO_4_. (**C**) Schematic representation of Pex14p showing the localization of *in vivo* phosphorylation sites. Phosphorylation at S6, S15, S76, T263, and S327 are reported here for the first time. MD, predicted membrane domain (according to Azevedo and Schliebs, 2006). Pex5p-, Pex7p-, and Pex13p-binding regions are depicted according to Azevedo and Schliebs (2006). Please note that alternative Pex14p-Pex13p binding sites have been proposed (Schell-Steven et al., 2005). While a second Pex14p interaction site has been mapped for Pex13p (Schell-Steven et al., 2005; Williams and Distel, 2006), an additional Pex13p binding site in Pex14p has not yet been reported. Coiled-coil (CC) domains were predicted using PredictProtein (Rost et al., 2004).

To identify and localize phosphorylation sites in native Pex14p, we used a multi-protease digestion approach followed by high resolution MS analysis. Affinity-purified native Pex14p was proteolytically cleaved in-solution with three different enzymes, *i*.*e*. Asp-N, Lys-C, or trypsin, to achieve full sequence coverage by LC-MS/MS analysis (**Supplementary Figure S1**). We identified 16 distinct Pex14p phosphosites with a localization probability of > 95% (**Table 1; Supplementary Table S4**), of which five (pS6, pS15, pS76, pT263, pS327) are reported here for the first time. As exemplarily shown in **Figure 1B** for the peptide carrying a phosphate moiety at serine 310 (pS310), localization of each phosphosite in Pex14p was confirmed by manual inspection and annotation of the corresponding MS/MS spectrum (**Supplementary Figure S2**). Interestingly, most of the phosphosites are clustered in the C-terminal region of Pex14p within the binding sites for the PTS1 and PTS2 receptor proteins Pex5p and Pex7p (**Figure 1C**). Moreover, our MS data reveal concurrent phosphorylation of Pex14p at S252/S254, S254/T263, S266/S268, T307/S310, T307/S313, S310/S313, and T307/S310/S313 (**Table 1**). In addition, four phosphosites are located in the N-terminal region within or close to the Pex5p- and Pex13p-binding sites (**Figure 1C**).

**Table 1.**
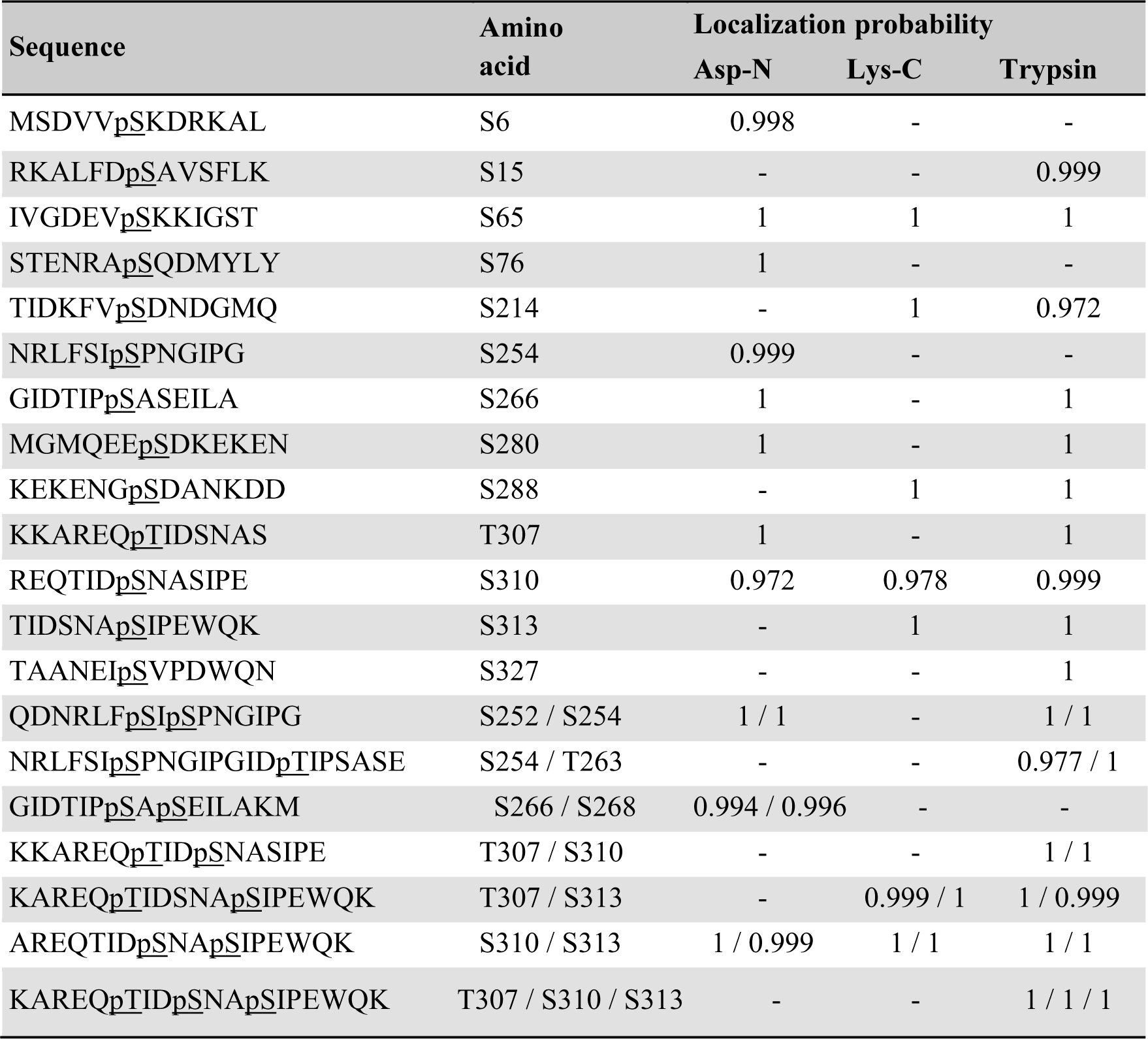
*In vivo* Pex14p phosphorylation sites identified by MS analyses of affinity-purified Pex14p^TPA^ following proteolytic digestion with Asp-N, Lys-C, or trypsin. Listed are phosphorylation sites of Pex14p phosphopeptides identified with a localization probability of ≥ 0.95 and a PEP value of < 0.01. The sequence shows the phosphorylated serine or threonine residues (<u>pS/T</u>) and up to six C- and N-terminally flanking amino acids. A comprehensive list of Pex14p (phospho)peptides is provided in **Supplementary Table S4**. The overall sequence coverage (excluding the methionine residue at position 1) was 100%; 97.4% were obtained with AspN, 93.3% with Lys-C, and 97.1% with trypsin (**Supplementary Figure S1**). n = 4 per protease.

### Expression of phosphomimetic Pex14p results in impaired growth on oleic acid and reduced peroxisomal protein import

To study the functional relevance of Pex14p phosphorylation, we generated yeast strains expressing phosphomimicking (exchange of serine and threonine to aspartate) and non-phosphorylatable (exchange to alanine) variants of wild-type Pex14p^TPA^ from its native chromosomal locus. We mutated all 16 Pex14p phosphosites to aspartate (16D mutant) and alanine (16A mutant).

Since the β-oxidation pathway in *S. cerevisiae* is restricted to peroxisomes, the functionality of peroxisomes can be determined by monitoring the growth of cells in medium containing oleic acid as carbon source. We therefore analyzed the ability of our mutant strains to grow on oleic acid in comparison to cells expressing wild-type Pex14p^TPA^ and *pex14Δ* cells. To exclude putative general growth defects, we also monitored growth in glucose-containing medium. As shown in **Figure 2A**, all strains showed no growth difference when glucose was present. However, in contrast to Pex14p^TPA^ wild-type cells and in line with published results (Albertini et al., 1997; Brocard et al., 1997), cells deficient in *PEX14* were unable to grow in oleic acid-containing medium. Interestingly, the phosphomimetic Pex14p^TPA^-16D mutant showed a severe growth defect in oleic acid medium that started to become evident approximately 25 h after shifting the cells from glucose to oleic acid while the non-phosphorylatable Pex14p^TPA^-16A mutant grew like Pex14p^TPA^ wild-type cells (**Figure 2A**). These data indicate that the phosphomimicking Pex14p mutant is significantly affected in its function, which results in the impairment of peroxisome metabolism and, thus, cellular growth on oleic acid.

Since Pex14p is a central component of the peroxisomal matrix protein import machinery, we next addressed the question whether the growth defect observed for the 16D mutant was caused by a deficiency in matrix protein import. To analyze the import efficiency of wild-type and mutant cells, we introduced a plasmid encoding for the synthetic peroxisomal matrix protein GFP C-terminally fused to the PTS1 sequence Ser-Lys-Leu (GFP-SKL) into our strains and a control strain expressing unmodified Pex14p and determined the intracellular localization of this artificial peroxisomal matrix protein. Strains expressing GFP-SKL were grown in oleic acid-containing medium and inspected by fluorescence microscopy for the cellular distribution of the fluorescence protein. In control cells, GFP-SKL was imported into peroxisomes as indicated by a punctate staining pattern characteristic for peroxisomes (Platta et al., 2004; **Figure 2B**). As expected, *pex14*Δ mutant cells show a diffuse fluorescence signal typical for a strain in which the PTS1-dependent protein import is defective and, as a consequence, peroxisomal matrix proteins are mislocalized to the cytosol (Distel et al., 1996). Strains expressing wild-type Pex14p^TPA^ or the Pex14p^TPA^-16A mutant were indistinguishable from control cells regarding their fluorescence pattern (**Figure 2B**). However, expression of GFP-SKL in the Pex14p^TPA^-16D mutant resulted in a punctate staining pattern like the one observed in control cells, but also a diffuse cytosolic fluorescence (**Figure 2B**), indicative for a decreased efficiency in peroxisomal matrix protein import.

In summary, our data suggest that the functionality of the peroxisomal protein import is impaired in cells expressing the phosphomimicking Pex14p^TPA^-16D mutant, leading to a growth defect on oleic acid medium.

### Impaired growth and reduced protein import can be attributed to phosphorylation of Pex14p at serine 266

To disclose if the phenotype of the Pex14p^TPA^-16D mutant can be attributed to a distinct phosphorylation site, we generated yeast strains expressing single Pex14p^TPA^ phosphosite mutants (i.e., single exchange of S or T to A and D) of all 16 phosphorylation sites identified in this work. Pex14p variants were expressed from native chromosomal location and the matrix protein import efficiency of the individual strains was analyzed using GFP-SKL and fluorescence microscopy (**Figure 3A**). Only the strain expressing the Pex14p^TPA^-S266D mutant exhibited enhanced cytosolic fluorescence of GFP-SKL as observed for the Pex14p^TPA^-16D mutant while the fluorescence pattern of the remaining phosphomimicking and of all non-phosphorylatable mutants resembled the pattern of wild-type cells (**Figure 3A**). In line with the data obtained for the Pex14p^TPA^-16D mutant, cells expressing Pex14p^TPA^-S266D are significantly impaired in growth on oleic acid-containing medium (**Figure 3B**). Immunoblot analysis of Pex14p^TPA^ wild-type, the S266A or S266D phosphosite mutant variant revealed comparable steady-state levels for Pex14p^TPA^ (**Supplementary Figure S3**), implying that the phenotype seen for cells expressing Pex14p^TPA^-S266D does not result from differences in Pex14p^TPA^ abundance.

**Figure 2.**
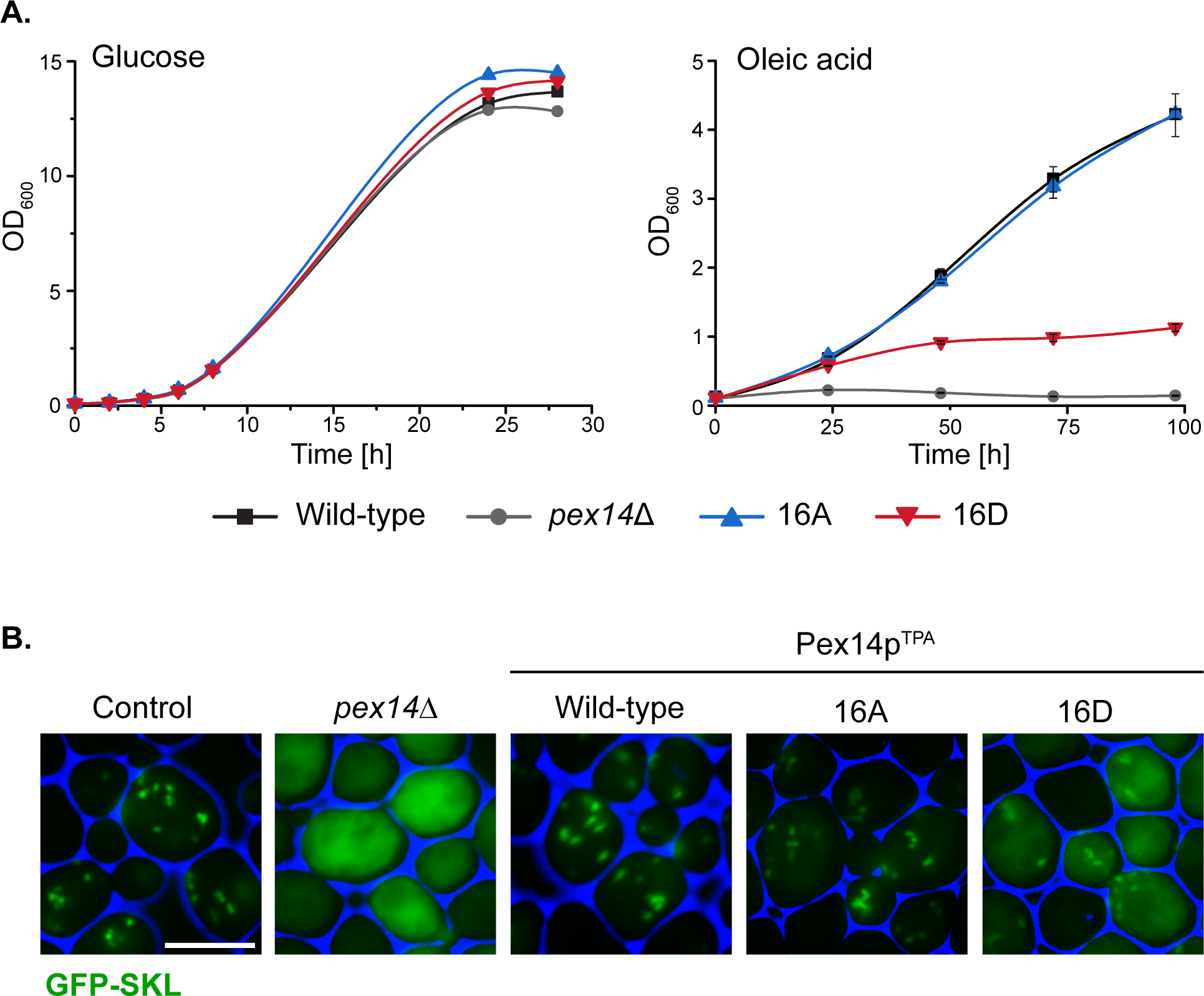
The Pex14p-16D mutant shows impaired growth on oleic acid and reduced peroxisomal import of GFP-SKL. (**A**) Strains were pre-cultured in medium containing 0.3% glucose and grown for 16 h at 30°C. Cells were shifted to either glucose (2%) or oleic acid medium, samples were taken at indicated time points, and the OD_600_ was determined. Growth in oleic acid medium was analyzed in triplicates. Error bars indicate standard deviation. (**B**) Representative fluorescence images of strains expressing native Pex14p (control) and mutant forms as well as plasmid-encoded GFP-SKL are shown. Cells were cultured in oleic acid medium as described in (**A**) and analyzed 16 h after the shift to oleic acid. The *pex14*Δ strain serves as control for a defect in peroxisomal protein import. Scale bar, 5 µm.

**Figure 3.**
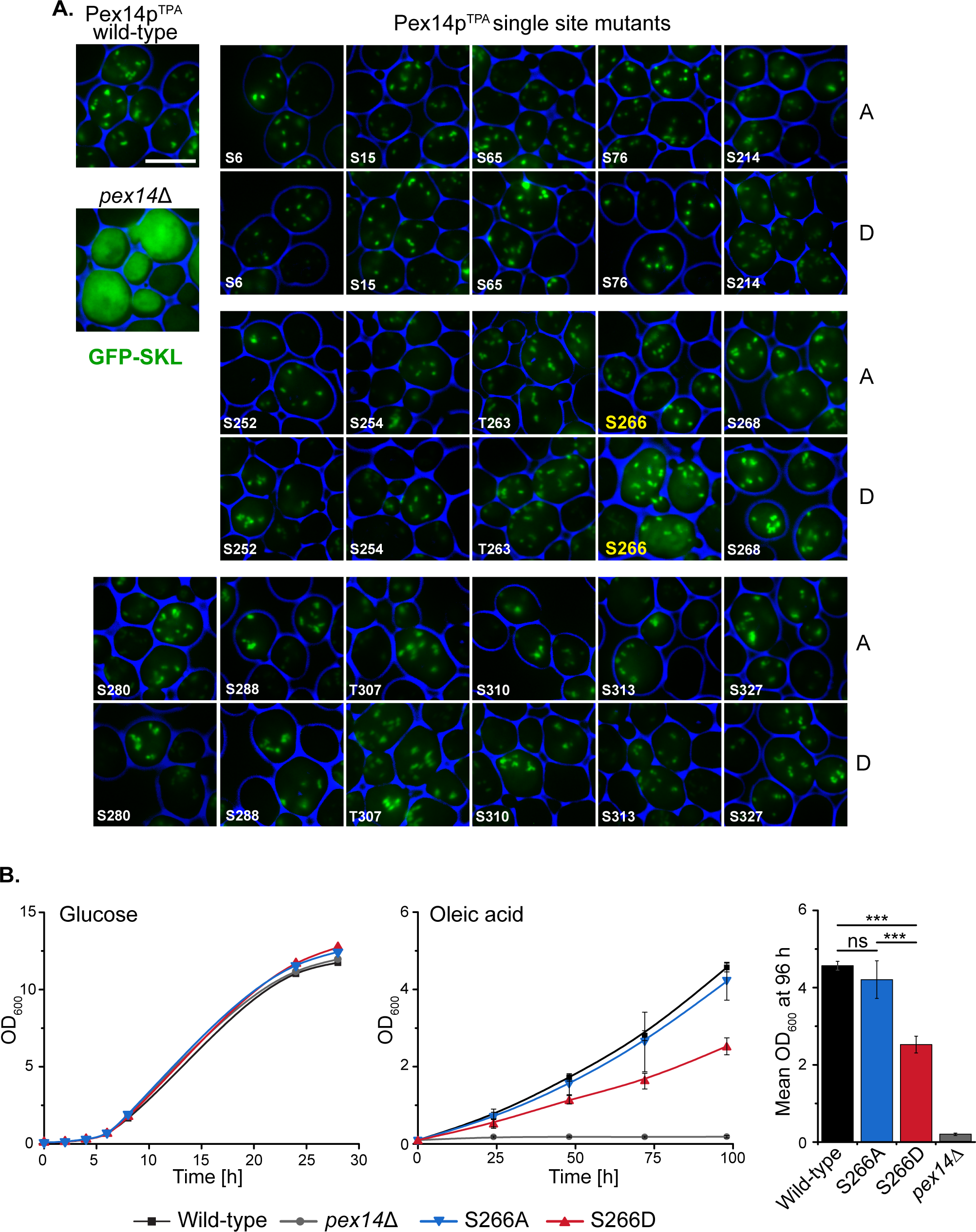
Deficiency in peroxisomal protein import and growth are caused by the phosphomimicking mutation at serine 266 of Pex14p. (**A**) Cells expressing Pex14p^TPA^ wild-type, single phosphorylatable or phosphomimicking (D) mutations as indicated and plasmid-encoded GFP-SKL were grown on oleic acid medium and analyzed as described in **Figure 2B**. *pex14*Δ cells serve as control for non-functional import of GFP-SKL. Scale bar, 5 µm. (**B**) Growth of strains in glucose- or oleic acid-containing medium was analyzed as described in **Figure 2A**. OD_600_ values of the 96-h time point for growth in oleic are visualized as bar chart (right plot). Error bars indicate standard deviation (n = 3); ***, p-value ≤ 0.001; ns, not significant.

Taken together, these data suggest that phosphorylation of Pex14p at S266 plays a crucial role for Pex14p function in peroxisomal protein import.

### Effect of Pex14p-S266 phosphosite mutants on the composition of peroxisomal import complexes

The impairment in growth and protein import observed for the Pex14p^TPA^-S266D mutant prompted us to analyze the capacity of Pex14p-S266 phosphosite mutants to associate with components of peroxisomal import complexes. Here, Pex14p phosphorylation may affect or regulate the binding to cargo-loaded receptor proteins and/or other proteins of peroxisomal import complexes. Protein complexes containing Pex14p wild-type or S266 phosphosite mutant forms were isolated from digitonin-solubilized membranes by affinity chromatography using the various TPA-tagged Pex14p proteins as baits. In line with previous work (Agne et al., 2003), wild-type Pex14p associates with its docking complex partner proteins Pex13p and Pex17p as well as Pex5p and Pex7p, as revealed by immunoblot analysis (**Figure 4A**, lane 6). Notably, Pex14p was isolated in similar amounts from cells expressing Pex14p^TPA^ wild-type or the mutant variants (**Figure 4A**, lanes 6 - 8) and equal amounts of Pex17p, Pex5p, and Pex7p were present in the respective Pex14p complexes. Only Pex13p appears to be less abundant in both the Pex14p-S266A and the -266D mutant complex compared to the wild-type.

**Figure 4.**
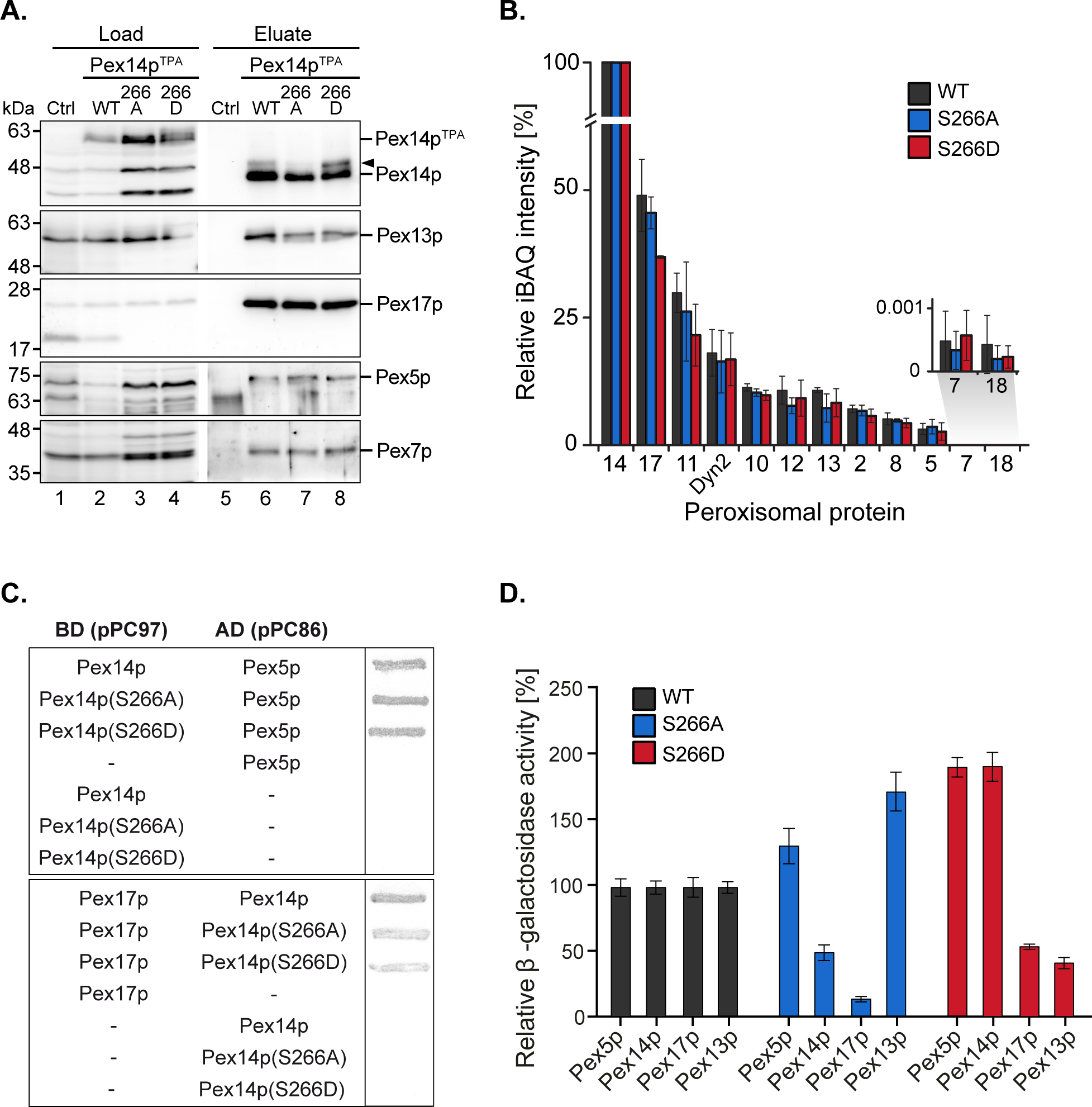
Composition of Pex14p-containing complexes. (**A**) *S. cerevisiae* control cells and cells with genomic integration of Pex14p^TPA^ wild-type and S266A and S266D phosphosite mutants were grown in oleic acid-containing medium for 16 h. Membrane protein complexes were solubilized from total cellular membranes using 1% (w/v) digitonin. Equal amounts of these protein mixtures containing either TPA-tagged Pex14p proteins or non-tagged Pex14p as control for unspecific binding were subjected to affinity chromatography with IgG-coupled Sepharose. Proteins were eluted by TEV protease cleavage. Proteins of the solubilized membrane fractions (load; 0.1% of total) and the eluate fractions (5% of total) were separated by SDS-PAGE followed by immunoblotting using the indicated antibodies. The arrowhead points to phosphorylated forms of Pex14p. Ctrl, control; WT, wild-type. (**B**) Eluates of the experiment shown in (**A**) and a second replicate were analyzed by LC-MS. Shown are mean iBAQ intensities and the standard deviation across both replicates. iBAQ intensities were normalized to the sum of all proteins identified in the respective LC-MS run and the values for Pex14p were set to 100%. iBAQ, intensity-based absolute quantification. (**C, D**) Interactions between distinct Pex14p variants and individual proteins required for peroxisomal protein import were analyzed using yeast two hybrid methods. Proteins were fused to either the binding or activation domain (BD, AD) of GAL4. The indicated plasmid combinations were co-transformed into the yeast two hybrid strain PCY2*pex14Δ*. Double transformants expressing the indicated combinations of Gal4p fusion proteins were (**C**) subjected to ß-galactosidase filter assay or (**D**) lysed and subjected to a liquid ß-galactosidase assay. The ß-galactosidase activity shown is the average of triplicate measurements for three independent transformants harboring each set of plasmids. Error bars denote standard error of the mean.

To obtain a more comprehensive picture of components of the peroxisomal import machineries that are associated with Pex14p complexes, the experiment was repeated and the eluates of both replicates were analyzed by high resolution LC-MS. This allowed for (i) the identification of a larger set of peroxisomal proteins associated with Pex14p, which for immunoblot analyses is limited by the availability of suitable antibodies, and (ii) the relative quantification of proteins in the Pex14p complexes (**Figure 4B; Supplementary Table S5**). In addition to the proteins detected by immunoblotting (**Figure 4A**), we identified in the eluates of Pex14p^TPA^ wild-type, -S266A and -266D affinity purifications all proteins that have previously been reported to form a Pex14p core complex: the RING finger complex (Pex2p, Pex10p, and Pex12p), the intraperoxisomal Pex8p, the dynein light chain protein Dyn2p, and Pex11p (Agne et al., 2003; Oeljeklaus et al., 2012; Chan et al., 2016) as well as the PTS2 co-receptor Pex18p, which together with Pex14p and Pex17p forms the peroxisomal PTS2 pore (Montilla-Martinez et al., 2015). However, significant differences in the abundance of individual peroxisomal proteins between the Pex14p variants were not observed, except for Pex17p, which appears to be slightly less abundant in Pex14p^TPA^-S266D complexes (**Figure 4B; Supplementary Table S5**). In sum, our data indicate that the phosphomimicking S266D mutation leads to slightly reduced levels of both Pex13p and Pex17p in Pex14p complexes at steady-state levels, which in the Pex14p-S266A mutant is only observed for Pex13p.

We next addressed the question if direct interactions between the Pex14p phosphosite mutants and its partner proteins in the docking complex as well as Pex5p are altered. Using yeast two-hybrid (Y2H) filter assays, we show that the Pex14p-S266A/D mutants interact with the PTS1 receptor Pex5p at wild-type level (**Figure 4C**, top), whereas the interaction of both the S266A and the S266D mutant with Pex17p was considerably decreased (**Figure 4C**, bottom). For Pex13p, we could not detect binding to any of the Pex14p variants (data not shown). The results of the filter assays were quantified by liquid Y2H assay (**Figure 4D**), which confirmed the reduced interaction of both Pex14p mutants with Pex17p. The Pex14p-S266A mutant further showed a deficiency in Pex14p self-association in the liquid Y2H assay (**Figure 4D**), an effect that does not result from varying expression levels of the different Pex14p variants (data not shown). Distinct from this, the Pex14p-S266D mutant exhibited a strongly reduced association to Pex13p, which was not observed for Pex14p-S266A (**Figure 4D**).

Taken together, our protein interaction data suggest that impaired growth and peroxisomal import of GFP-SKL may be a consequence of a reduction in the association of the phosphomimicking Pex14p-S266D mutant to its partner proteins Pex13p and Pex17p.

### The Pex14p^TPA^-266D mutant affects the subcellular distribution of peroxisomal citrate synthase

The decreased peroxisomal import that we observed for the Pex14p^TPA^-S266D mutant was revealed using the artificial cargo protein GFP-SKL. To view a more physiological situation, we analyzed the effect of Pex14p phosphosite mutants on the import of individual peroxisomal matrix proteins. To this end, we generated yeast strains each expressing either Pex14p^TPA^ wild-type, the 266D or S266A variant from its native locus, a selected peroxisomal matrix protein N-terminally tagged with GFP, and Pex3p-mCherry as peroxisomal marker protein. GFP-tagged proteins were expressed under the control of a constitutive *SpNOP1* promoter to facilitate the detection of low abundance proteins and proteins that are only expressed under specific conditions (Yifrach et al., 2016; Yofe et al., 2016; Dahan et al., 2017). The correct targeting of GFP-tagged peroxisomal proteins including matrix proteins with an N-terminal PTS2 to peroxisomes has been shown before (Yofe et al., 2016). Cells were grown in glucose-containing medium and analyzed by fluorescence microscopy using a high-content screening platform (Breker et al., 2013). The subcellular localization of 23 peroxisomal matrix proteins that are imported into peroxisomes via different import pathways (e.g. PTS1 or PTS2 pathway, piggyback import) was assessed manually. Our analysis revealed that the cellular distribution of the majority of these proteins remained unaltered in the Pex14p^TPA^-S266 mutants (**Figure 5A; Supplementary Figure S4**). In contrast, we noticed differences in the localization of GFP-Aat2p, an aspartate aminotransferase (**Figure 5B**), and GFP-Cit2p, the peroxisomal isoform of citrate synthase involved in the glyoxylate cycle (**Figure 5C**). In Pex14^TPA^ wild-type cells, GFP-Aat2p is localized to peroxisomes as indicated by a punctate pattern that is congruent to the red fluorescence pattern of Pex3p-mCherry. In addition, weak cytosolic fluorescence is seen, which is much stronger in the 266A and the 266D mutant (**Figure 5B**). However, both the non-phosphorylatable and the phosphomimicking Pex14p^TPA^ mutant show the same phenotype, suggesting that the observed effect does not depend on Pex14p phosphorylation at S266.

In accordance with a dual localization of Cit2p in peroxisomes and the cytosol (Nakatsukasa et al., 2015), GFP-Cit2p accumulates in peroxisomal puncta and shows a substantial cytosolic distribution in Pex14p^TPA^ wild-type and -S266A cells (**Figure 5C**). The same staining pattern is also observed for Pex14p^TPA^-S266D cells, but with an increased fluorescence signal in the cytosol. This indicates that phosphorylation of Pex14p at S266 exerts an adverse effect on the peroxisomal import of GFP-Cit2p under the experimental conditions employed here.

Cit2p catalyzes the first reaction of the glyoxylate cycle, which is required for the production of carbohydrates when yeast cells are grown on non-fermentative carbon sources such as acetate or fatty acids. Since in the fluorescence microcopy screen GFP-Cit2p was expressed under the control of the constitutive *SpNOP1* promoter and cells were grown in glucose, which promotes fermentative cell growth in *S. cerevisiae*, we next analyzed the subcellular localization of GFP-Cit2p expressed under regulation of its native promoter in cells grown on oleic acid. Fluorescence microscopy analysis of seamless GFP-Cit2p cells (Yofe et al., 2016) expressing the different Pex14p^TPA^ variants confirmed the results of the initial screen for oleic acid-grown cells: irrespective of the Pex14p^TPA^ variant, seamless GFP-Cit2p-expressing cells show a punctate staining pattern and diffuse cytosolic fluorescence, which is much stronger in Pex14p^TPA^-S266D cells (**Figure 5D**). Quantification of the cytosolic fluorescence revealed a significantly higher cytosolic fluorescence in S266D cells compared to wild-type and S266A cells, which was not the case between wild-type and S266A mutant cells (**Figure 5E**). This effect was not a result of different GFP-Cit2p or Pex14p^TPA^ expression levels (**Supplementary Figure S5A**). Consistent with the data obtained in the initial screen, the subcellular distribution of GFP-Mdh3p, a known PTS1 protein, remained unaltered (**Supplementary Figures S5B** and **S5C**).

To corroborate our fluorescence microscopy data, we analyzed the subcellular distribution of GFP-Cit2p in Pex14p^TPA^ wild-type, -S266D and -S266A cells in sedimentation assays. Postnuclear supernatants of cells grown in oleic acid were separated into a cytosolic fraction and an organellar pellet by centrifugation through a sucrose layer. Fractions were analyzed for the presence of GFP-Cit2p, Pex14p^TPA^ as well as peroxisomal, mitochondrial and cytosolic marker proteins by immunoblotting (**Figure 5F**). The detection of the cytosolic marker protein 3-phosphoglycerate kinase (Pgk1p) in the soluble fraction of all strains (lanes 2, 8, 11) and mitochondrial porin (Por1p) detection exclusively in the organellar pellets (lanes 3, 9, 12) confirm proper cell fractionation. Pex14p^TPA^ was consistently present only in the postnuclear supernatant and the organellar pellet, demonstrating that all three Pex14p^TPA^ variants are effectively targeted to peroxisomes without any observable mislocalization to the cytosol. Consistent with a peroxisomal localization and our fluorescence microscopy data, Cta1p and Pcs60p, two PTS1 proteins, were almost entirely present in the organellar pellet with only very minor amounts detected in the cytosolic fraction, irrespective of the Pex14p^TPA^ variant expressed (**Supplementary Figure S4; Figure 5F**). In *pex14Δ* cells, which served as control, these peroxisomal matrix proteins were mislocalized to the cytosol (lane 5). Our immunoblot data further confirm the dual localization of GFP-Cit2p: in Pex14p^TPA^ wild-type and -S266A cells, GFP-Cit2p was roughly equally split between cytosolic and organellar fraction (lanes 2/3, 8/9). However, in Pex14p^TPA^-S266D cells, GFP-Cit2p was clearly shifted to the cytosolic fraction along with a strong reduction in the organellar pellet fraction (lanes 11/12), which is consistent with the fluorescence microscopy data **(Figures 5D and 5E)**. Quantification of immunoblot signal intensities for GFP-Cit2p, Pex14p^TPA^ and Cta1p further corroborates our observation of a reduced peroxisomal import of GFP-Cit2p in the Pex14p^TPA^-S266D mutant.

In conclusion, our data point to a mechanism that controls the peroxisomal import of Cit2p via the state of Pex14p-S266 phosphorylation.

### The Pex14p^TPA^-S266D phosphosite mutant has a growth advantage on ethanol compared to the S266A mutant

Cit2p catalyzes the condensation of acetyl-CoA with oxaloacetate to form citrate. When *S. cerevisiae* is grown on oleic acid, the fatty acid is degraded via the β-oxidation pathway in peroxisomes and its end product acetyl-CoA is fed into the glyoxylate cycle, which requires Cit2p activity in peroxisomes. An alternative carbon source that can be utilized by *S. cerevisiae* is ethanol, which is converted to acetyl-CoA via a series of reactions in the cytosol. Thus, when yeast is grown on ethanol, cytosolic citrate synthase activity is required to feed acetyl-CoA into the glyoxylate cycle. This prompted us to ask if growth of the S266 phosphosite mutants was different on ethanol compared to oleic acid. Indeed, when Pex14p^TPA^ wild-type, -S266A and -S266D cells were grown on ethanol, the S266D mutant grew like wild-type cells, whereas the S266A mutant was significantly impaired in growth (**Figure 6**), indicating that S266D cells, in which Cit2p is retained in the cytosol, have a clear advantage over S266A cells under ethanol growth conditions.

**Figure 5.**
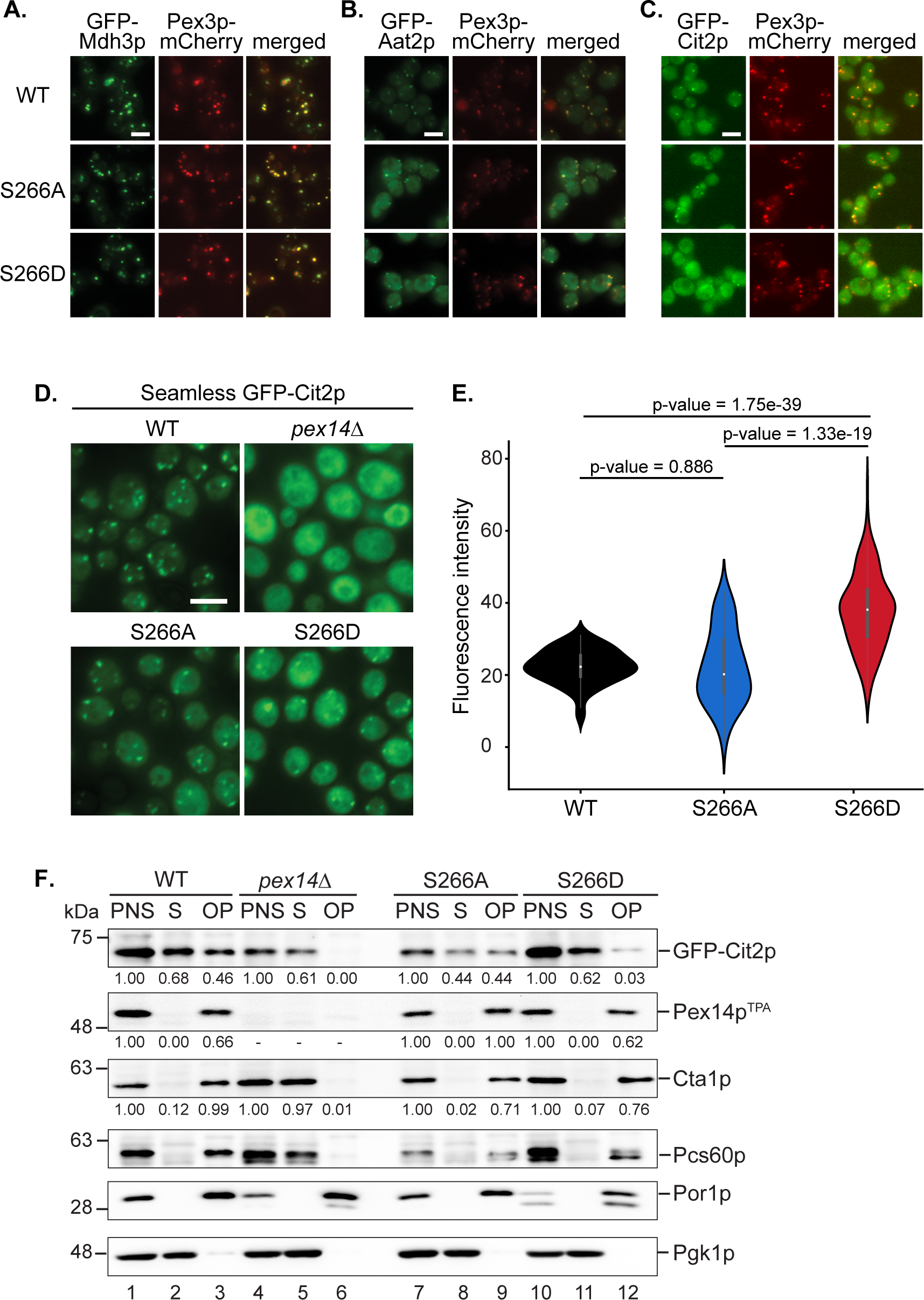
The phosphomimicking Pex14p-S266D mutant affects peroxisomal import of Cit2p. (**A-C**) The subcellular distribution of GFP-tagged peroxisomal matrix proteins was analyzed by fluorescence microscopy in strains expressing Pex14p^TPA^ wild-type (WT), the S266A or S266D mutant and Pex3p-mCherry as peroxisomal marker protein. GFP-tagged proteins were expressed under control of the constitutive *SpNop1* promoter and cells were grown in glucose-containing medium. Shown are representative images for GFP-Mdh3p (**A**), GFP-Aat2p (**B**), and GFP-Cit2p (**C**). See also **Supplementary Figure S4**. Scale bars, 5 µm. (**D**) Cells expressing the different Pex14p^TPA^ variants and GFP-Cit2p under control of the native promoter (‘seamless’) were grown in oleic acid medium as described in **Figure 2A** and analyzed by fluorescence microscopy 16 h after the shift. *pex14*Δ cells serve as control for a defect in peroxisomal matrix protein import. Scale bar, 5 µm. (**E**) Quantification of the cytosolic fluorescence in Pex14p^TPA^-WT, -S266A and -S266D cells grown in oleic acid as exemplarily shown in (D). p-values were determined using the Welch’s test (n = 73 for WT, n = 69 for S266A, n = 174 for S266D). (**F**) Seamless GFP-Cit2p cells expressing Pex14p^TPA^ variants as indicated or lacking *PEX14* were grown in oleic acid and used to prepare postnuclear supernatants (PNS), which were subsequently separated into a supernatant (S, cytosolic fraction) and an organellar pellet (OP). Equal portions of PNS, S and OP fractions were analyzed by immunoblotting using antibodies recognizing the indicated proteins. Numbers below immunoblot signals indicate relative signal intensities normalized to the signal detected in the corresponding PNS.

**Figure 6.**
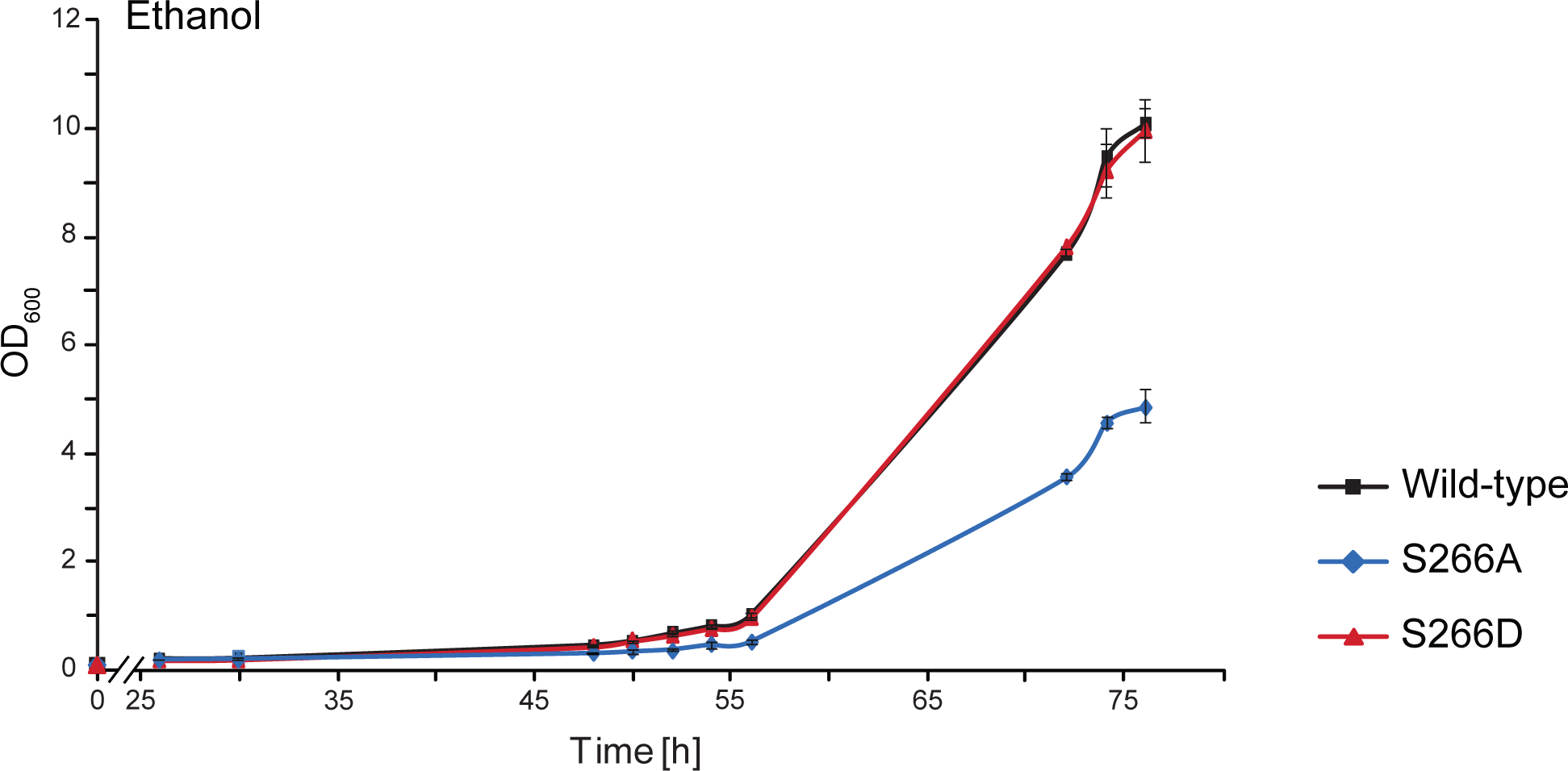
Growth of yeast cells expressing different Pex14p^TPA^ variants on ethanol. Pex14p^TPA^ wild-type, S266A or S266D cells were pre-cultured in medium containing 0.3% glucose, grown for 16 h at 30°C and shifted to ethanol (2%) medium. Samples were taken at indicated time points and the OD_600_ was determined. Error bars indicate standard deviation (n = 3).

Taken together, our data suggest that reduced peroxisomal import of Cit2p is beneficial for the cell to grow on ethanol and that the subcellular localization of Cit2p may likely be regulated in response to the cellular location of acetyl-CoA production in a Pex14p-S266 phosphorylation-dependent manner.

## Discussion

Pex14p is essential for peroxisomal matrix protein import. Together with Pex13p, and Pex17p in yeast, it forms the peroxisomal receptor docking complex (Agne et al., 2003). In addition, Pex14p is part of both the PTS1 and the PTS2 import pore (Meinecke et al., 2010; Montilla-Martinez et al., 2015). While the key players of the peroxisomal matrix protein import machinery in yeast have been identified (Walter and Erdmann, 2019), the molecular mechanisms controlling this process and the proteins involved are still largely unknown. Here, we established an *in vivo* phosphorylation map of Pex14p of *S. cerevisiae* comprising 16 phosphosites that are mainly localized in the Pex5p- and Pex7p-binding regions of Pex14p (**Figure 1C**). Of note, the phosphorylatable residues S15, S65, and S266 are conserved in human, mouse, and rat.

Phosphorylation of Pex14p has also been reported for the methylotrophic yeasts *Hansenula polymorpha* (Komori et al., 1999; Komori and Veenhuis, 2000; Tanaka et al., 2013) and *Pichia pastoris* (Johnson et al., 2001; Farre et al., 2008). *Hp*Pex14p phosphorylation has been discussed to be involved in pexophagy. However, direct experimental evidence supporting this assumption is still missing (de Vries et al., 2006). Exchange of a threonine and serine residue in *Hp*Pex14p, known to be phosphorylated *in vivo*, against non-phosphorylatable alanine had no effect on pexophagy or peroxisomal protein import under the experimental conditions chosen (Tanaka et al., 2013). In *S. cerevisiae*, however, Pex14p is not involved in pexophagy (Motley et al., 2012b), instead, Pex3p is the key molecule for the degradation of peroxisomes (Motley et al., 2012a;b).

To gain insight into the functional role of Pex14p phosphorylation of baker’s yeast, we generated yeast cells expressing phosphomimicking and non-phosphorylatable mutants of all 16 phosphosites. For the phosphomimicking 16D mutant, we observed a growth defect and impaired import of GFP-SKL into peroxisomes (**Figure 2**), which points to a role of Pex14p phosphorylation in modulating the import of peroxisomal matrix proteins. However, we cannot rule out that the simultaneous exchange of 16 serine/threonine residues to aspartate, which is associated with the introduction of additional negative charges in particular in the C-terminus of the protein, has adverse effects on the properties of Pex14p interfering with efficient matrix protein import and, consequently, growth on oleic acid.

To exclude such an effect, we analyzed single mutants of all 16 phosphosites identified in Pex14p of *S. cerevisiae*. It turned out that phosphorylation at S266 plays a crucial role in matrix protein import, as the phosphomimetic version S266D displays reduced import for GFP-SKL and impaired growth on oleic acid (**Figure 3**). However, the decrease in growth was more pronounced for the 16D mutant (**Figure 2A**), most likely in consequence of charge-induced alterations in the protein structure or its ability to form intra- or intermolecular protein interactions as discussed above. Thus, the 16D variant of Pex14p is most likely more compromised in its overall function(s) than the single site S266D mutant, which provides a reasonable explanation for the differences in the growth defect (**Figures 2A and 3B**).

Wild-type Pex14p is known to interact with the membrane proteins Pex13p and Pex17p as well as the PTS1-receptor Pex5p (Albertini et al., 1997; Huhse et al., 1998). In line with this finding, the S266D mutant of Pex14p is still able to form the receptor-docking complex and to associate with Pex5p. However, while Pex5p-binding of both Pex14p-S266A and -S266D is not impaired, both variants display reduced binding of Pex17p (**Figure 4**). But why is there just an import defect seen for the S266D mutant? The molecular reason behind this may be an additional reduced binding capacity of this Pex14p variant to Pex13p. While the S266A mutant interacts with this docking complex constituent like wild-type Pex14p, binding of Pex13p is reduced to < 50% of wild-type levels for the S266D mutant (**Figure 4D**). It is known that Pex14p interacts with the SH3 domain of Pex13p via a P-x-x-P motif in its N-terminus (amino acid residues 87 - 92, Girzalsky et al., 1999; **Figure 1C**). Mutation of the proline residues in this motif to alanine abolishes the interaction with the Pex13p-SH3 domain. However, the Pex14p A-x-x-A mutant is still able to bind to Pex13p, pointing to the existence of alternative Pex13p-Pex14p binding sites (Schell-Steven et al., 2005). For Pex13p, a second Pex14p binding site has been identified (Schell-Steven et al., 2005; Williams and Distel, 2006), but a second binding site for Pex13p in Pex14p has not yet been reported. Thus, it is tempting to speculate that S266 may be part of this potential second Pex13p interaction site and that the Pex14p-Pex13p association at this site may be modulated via the state of S266 phosphorylation.

Taken together, our protein interaction data suggest that the combined effect of reduced association with both Pex13p and Pex17p causes the observed reduced peroxisomal import of GFP-SKL in cells expressing Pex14p-S266D. This is in agreement with the function of the Pex13p/Pex14p subcomplex as docking platform for receptor-cargo complexes (Elgersma et al., 1996; Erdmann and Blobel, 1996; Gould et al., 1996) and the proposed role of Pex17p to increase the efficiency of the docking event (Chan et al., 2016).

Most interestingly, among 23 native matrix proteins tested, GFP-Cit2p was the only protein that displayed an altered cellular distribution depending on the Pex14p^TPA^-S266 phosphosite mutant, exhibiting an increased cytosolic localization in the phosphomimicking S266D mutant (**Figure 5A-C; Supplementary Figure S4**). This effect was most evident in cells expressing GFP-Cit2p at endogenous Cit2p levels grown on oleic acid (**Figure 5D**), a metabolic condition that requires the activity of the glyoxylate cycle for the production of essential carbohydrates (Kunze et al., 2006). Cit2p catalyzes the condensation of acetyl-CoA with oxaloacetate to form citrate. Subsequent reactions of the glyoxylate cycle comprise the isomerization of citrate to isocitrate, cleavage of isocitrate to succinate (the product of the glyoxylate, serving as precursor for carbohydrates via gluconeogenesis) and glyoxylate, condensation of glyoxylate and acetyl-CoA yielding malate, and oxidation of malate to oxaloacetate, required to maintain metabolic flux through the glyoxylate cycle. In *S. cerevisiae*, the reactions are partitioned between peroxisomes and the cytosol (Minard and McAlister-Henn, 1991; Chaves et al., 1997; Kunze et al., 2002; Regev-Rudzki et al., 2005). Cit2p has been shown to be active in both compartments (Lee et al., 2011; Nakatsukasa et al., 2015). Cellular abundance of Cit2p is controlled on both transcriptional and post-translational level. Expression of the *CIT2* gene is controlled via the retrograde response pathway, which mediates communication from mitochondria to the nucleus in response to mitochondrial integrity and metabolic needs of the cell (Liao et al., 1991; Liu and Butow, 2006). Post-translationally, Cit2p is ubiquitinated by the SCF-Ucc1p ubiquitin ligase complexes and proteasomally degraded in glucose-grown cells, when the activity of the glyoxylate cycle is not required. In contrast, in cells grown on the acetyl-CoA precursor acetate, *UCC1* expression is downregulated, which prevents Cit2p degradation, and, thus, the glyoxylate cycle is active (Nakatsukasa et al., 2015).The docking complex at the peroxisomal membrane makes for an excellent checkpoint to control the import of matrix proteins via its central component Pex14p. Here, we propose that phosphorylation of Pex14p at S266 controls the distribution of Cit2p between peroxisomes and the cytosol by reducing the efficiency of its import into peroxisomes to adjust Cit2p localization to the location of acetyl-CoA production: Cit2p is shifted to peroxisomes when acetyl-CoA is generated from oleic acid via fatty acid β-oxidation and to the cytosol when acetyl-CoA is generated from ethanol in the cytosol. This adds a further level of Cit2p regulation. In line with this, the Pex14p-S266D mutant showed a growth advantage over the S266A mutant when cells were cultured with ethanol **(Figure 6)**, whereas the opposite growth behavior was observed on oleic acid (**Figure 2B**).

From the mechanistic point of view, it is striking that the peroxisomal import of only a single protein is affected in the Pex14p-S266D mutant, while the majority of peroxisomal matrix proteins are imported like in wild-type cells. Future studies will be directed at deciphering the molecular details underlying this regulatory mechanism. However, considering that S266 is located within the Pex5p-binding region of Pex14p, it is conceivable that the affinity of Cit2p-loaded Pex5p to Pex14p-S266D at the peroxisomal membrane is reduced. Cargo proteins differ in many characteristics (e.g. mass, shape/structure), and some proteins are even imported as oligomers, which has an impact on the overall structure of Pex5p-cargo complexes with different structures for different cargoes. As a consequence, Pex5p-cargo complexes may differ in their binding affinity to Pex14p in the peroxisomal docking complex, which in the Pex14p-S266D mutant may be more reduced for the Pex5p-Cit2p complex than for other Pex5p-cargo complexes. Moreover, it has been shown that peroxisomal targeting sequences differ in strength. Pex5p-cargo formation does not only rely on the C-terminal PTS1 tripeptide of the cargo protein but also on adjacent amino acid residues, with arginine at position -2 and arginine or lysine at -1 favoring binding of the extended PTS1 to Pex5p in *S. cerevisiae* (Lametschwandtner et al., 1998). The extended PTS1 of Cit2p features isoleucine at -2 and glutamic acid at -1 (ELVKNIE-SKL), which points to a weaker PTS1. Although this remains speculative, combined with a potentially reduced binding of the Pex5p-Cit2p complex to Pex14p-S266D, weaker binding of Cit2p to Pex5p in a context of other cargo proteins competing for Pex5p may result in a reduced efficiency of Cit2p translocation into the peroxisome. Furthermore, since Pex14p is part of the dynamic import pore, it may directly contribute to cargo translocation, which may be regulated by phosphorylation in a cargo-specific manner To conclude, we propose that the phosphorylation-mediated modulation of Cit2p import represents a mechanism that participates in fine-tuning the carbohydrate metabolism according to the nutritional demands of the cell. Taking our observations to a higher level, we envision that the finding that phosphorylation of a single residue in a peroxisomal biogenesis factor, such as Pex14p-S266 in our study, affects peroxisomal import of a specific protein is an intriguing new concept for the modulation and precise regulation of metabolic processes in a cell.

## Abbreviations

PTS: peroxisomal targeting signal;
LC-MS: liquid chromatography-mass spectrometry;
TEV: tobacco etch virus;
TPA: TEV protease cleavage site-Protein A

## Data availability statement

The datasets generated for this study are available on request to the corresponding author.

## Author contributions

AS performed phosphoproteomics, mass spectrometric and biochemical analyses as well as fluorescence microscopy. AS, RM, TH, and IS generated plasmids and yeast strains. RM, SGM and AF performed fluorescence microscopy analyses and WWDM quantified fluorescence microscopy data. RM conducted biochemical assays. TH performed yeast two hybrid assays. All authors analyzed data and interpreted experiments. BW conceived the project and individual experiments were designed by BW, SO, RE, WG, MS, and EZ. SO and BW wrote the manuscript together with WG and with input of other authors.

## Funding

This research was funded by the Deutsche Forschungsgemeinschaft (DFG, German Research Foundation) – FOR 1905 to BW and RE, Project-ID 278002225 – RTG 2202 and Project-ID 403222702 – SFB 1381 to BW, the Excellence Strategy (CIBSS – EXC-2189 – Project-ID 390939984 to BW), and the Excellence Initiative of the German Federal & State Governments (EXC 294, BIOSS to BW). This work was supported by a grant from the European Union Marie Curie Initial Training Networks (ITN) program PerICo (Grant Agreement Number 812968) to BW and RE. Work in the Schuldiner lab is supported by an ERC CoG Peroxisystem 646604. MS is an incumbent of the Dr. Gilbert Omenn and Martha Darling Professorial Chair in Molecular Genetics.

## Acknowledgements

We thank the staff of the Life Imaging Center (LIC) at the Center for Biological Systems Analysis (ZBSA) of the Albert-Ludwigs-University Freiburg for help with their confocal microscopy resources and the excellent support in imaging and analysis and Chris Meisinger for scientific discussion. The anti-Cta1p antibody was a kind gift of Andreas Hartig (University of Vienna, Austria). Work included in this study has also been performed in partial fulfillment of the doctoral theses of AS, RM, and TH.

## Supplementary Material

**Supplementary Figure S1**. Sequence coverage of Pex14p.

**Supplementary Figure S2**. Annotated MS/MS spectra of Pex14p peptide ions comprising the phosphorylation site(s) indicated in the figure.

**Supplementary Figure S3**. Cells expressing GFP-SKL and Pex14p^TPA^ wild-type, S266A or S266D exhibit comparable Pex14p^TPA^ steady-state levels.

**Supplementary Figure S4**. Fluorescence microscopy analysis of GFP-tagged peroxisomal matrix proteins in cells expressing Pex14p^TPA^ wild-type, the S266A or S266D mutant.

**Supplementary Figure S5**. Validation of data obtained in fluorescence microscopy experiments of GFP-Cit2p and GFP-Mdh3p cells.

**Supplementary Table S1**. Yeast strains used in this study.

**Supplementary Table S2**. Primers used in this study.

**Supplementary Table S3**. Plasmids and cloning strategies used in this study.

**Supplementary Table S4**. Pex14p (phospho)peptides identified by LC-MS in native Pex14p affinity-purified from crude membrane fractions of *S. cerevisiae*.

**Supplementary Table S5**. Results of LC-MS analyses of Pex14p complexes affinity-purified from cells expressing Pex14p^TPA^ wild-type or S266A and S266D phosphosite mutants thereof.

## Conflict of interest

The authors declare that the submitted work was carried out without any personal, professional or financial relationships that could be construed as a conflict of interest.

---

**Supplementary Figure S1:**
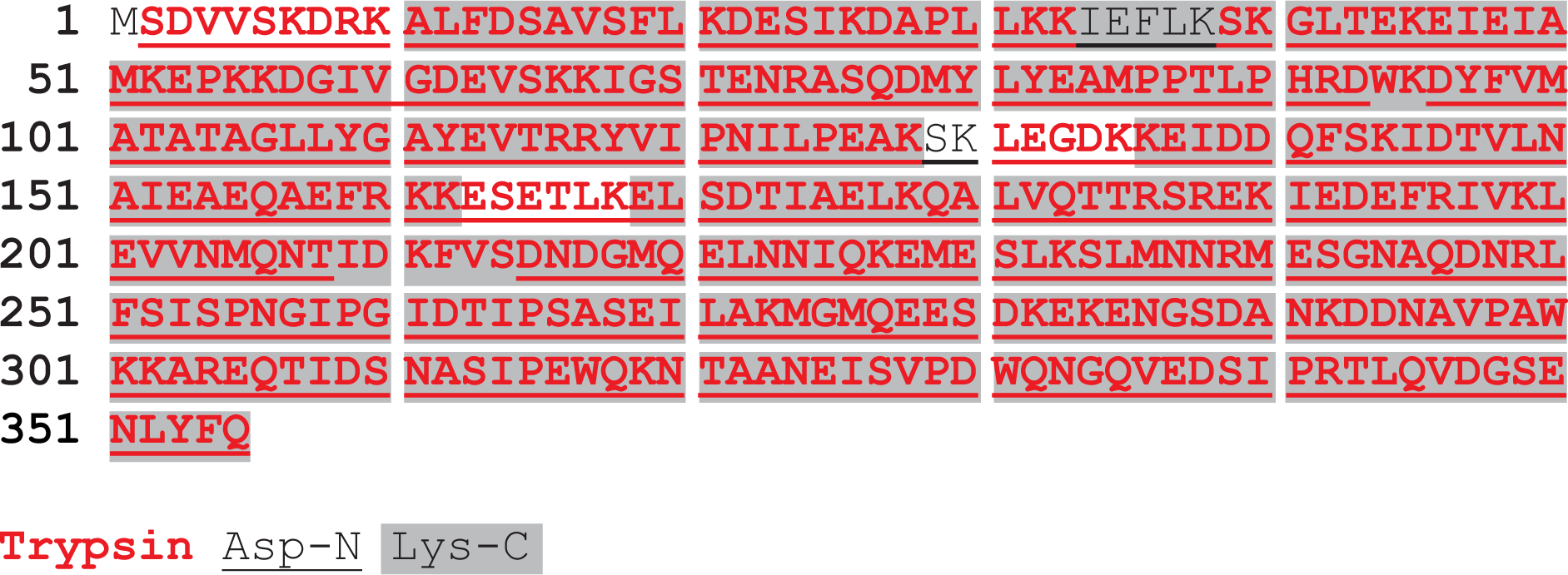
Sequence coverage of Pex14p. Pex14p was affinity-purified from crude membranes, proteolytically digested in solution with trypsin, Asp-N, or Lys-C, and subsequently analyzed by LC-MS. Tryptic peptides are shown in bold red, Asp-N peptides are underlined, and Lys-C peptides are highlighted in grey. Note that the sequence shown contains at the C-terminus the residue of the TPA tag (RTLQVDGSENLYFQ) that remained after TEV cleavages.

**Supplementary Figure S3:**
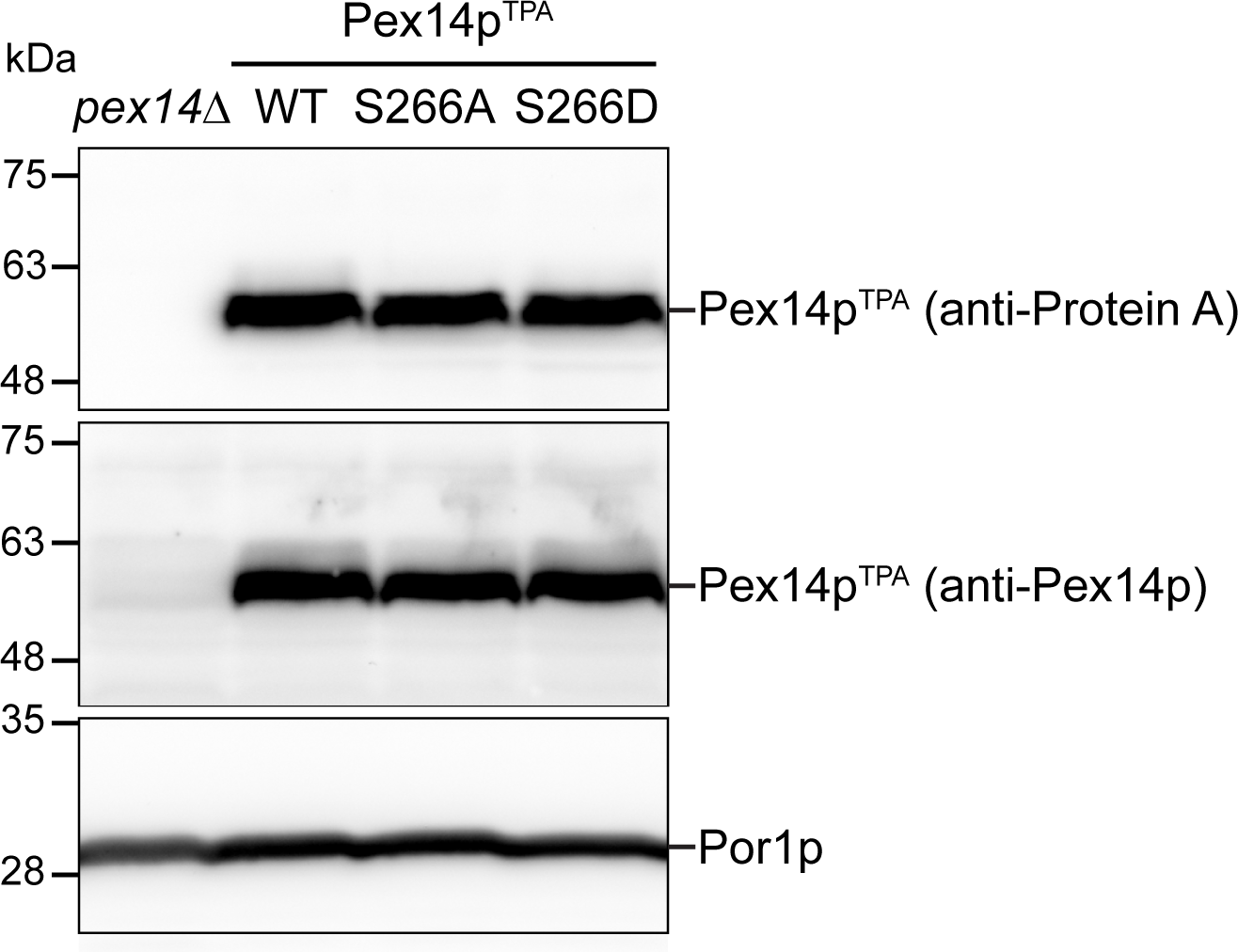
Cells expressing GFP-SKL and Pex14p^TPA^ wild-type, S266A or S266D exhibit comparable Pex14p^TPA^ steady-state levels. Whole cell lysates obtained from cell expressing GFP-SKL and the indicated Pex14p^TPA^ variants or lacking *PEX14* (*pex14*Δ) were analyzed by immunoblotting using antisera recognizing Protein A, Pex14p, and the mitochondrial protein Por1p (loading control). WT, wild-type.

**Supplementary Figure S4:**
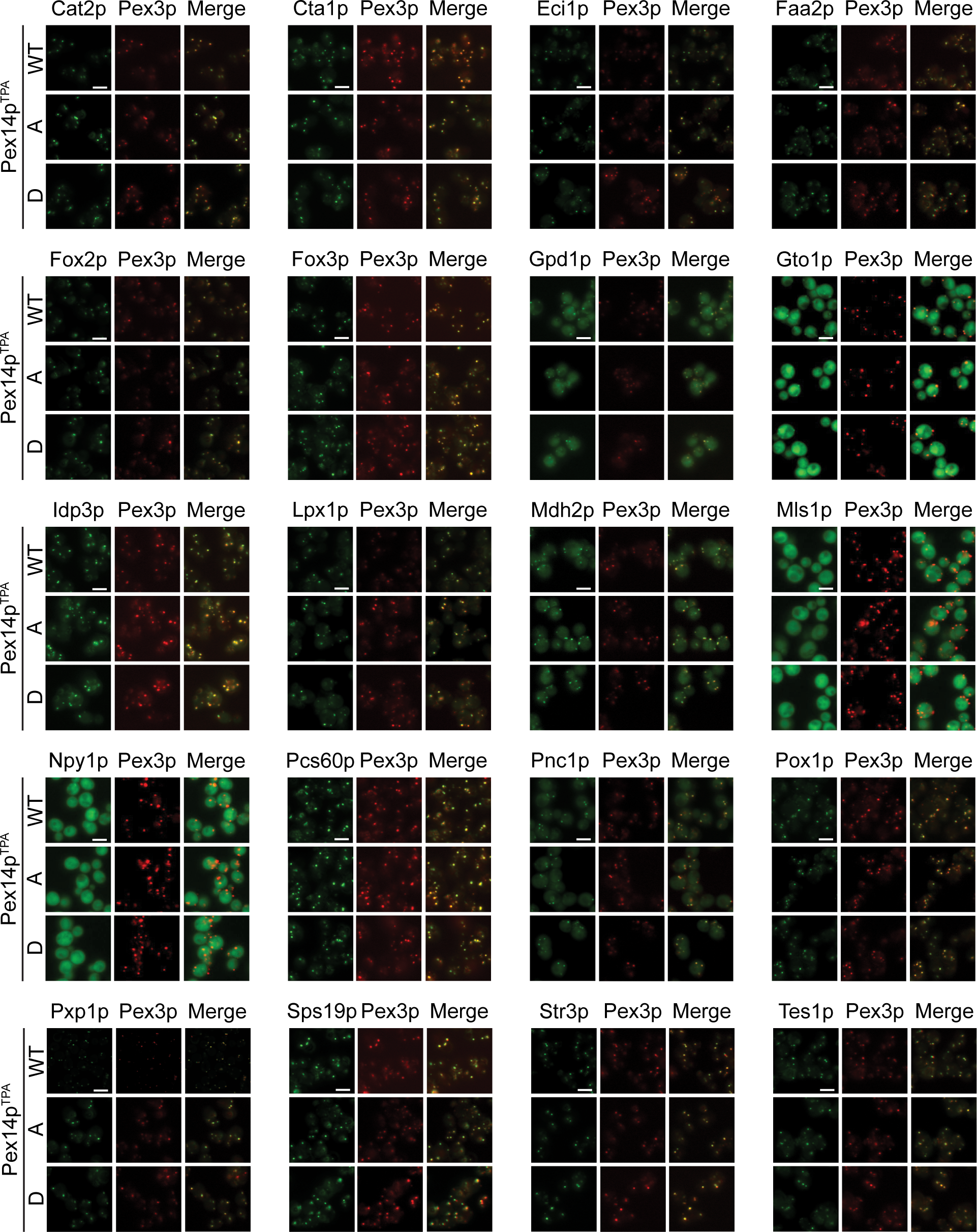
Fluorescence microscopy analysis of GFP-tagged peroxisomal matrix proteins in cells expressing Pex14p^TPA^ wild-type, the S266A or S266D mutant. Same experiment as described for Figure 5A-C. Shown are representative images for the indicated proteins. WT, wild-type; A/D, S266A/D mutant. Scale bars, 5 µm.

**Supplementary Figure S5:**
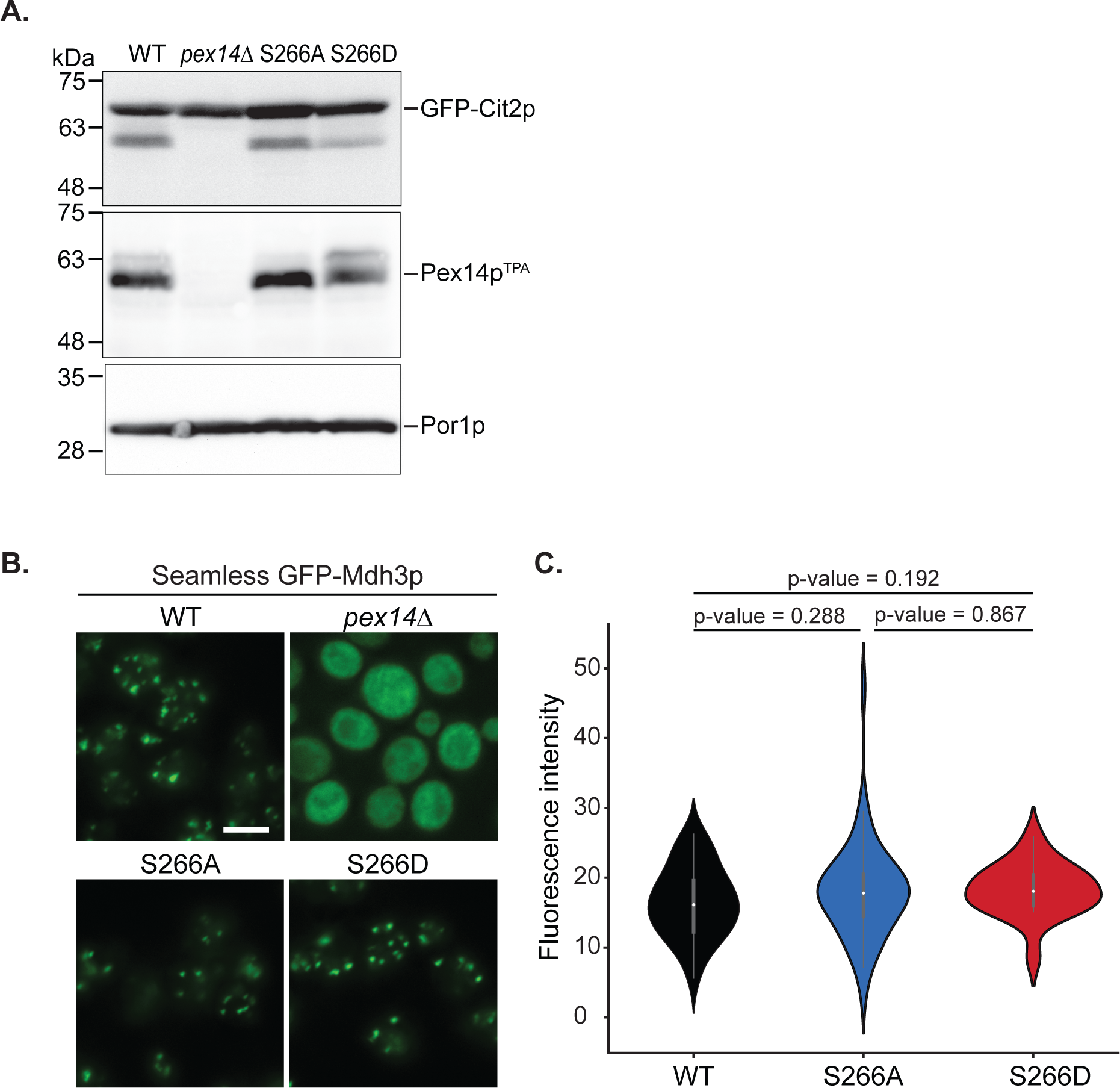
Validation of data obtained in fluorescence microscopy experiments of GFP-Cit2p and GFP-Mdh3p cells. (**A**) Whole cell lysates prepared from cells of the same cultures that were used for the fluorescence microscopy analysis shown in Figure 5D were analyzed by immunoblotting using antisera recognizing GFP, Pex14p, and the mitochondrial protein Por1p, serving as loading control. WT, wild-type. (**B**) Same experiment as shown in Figure 5D using cells expressing the different Pex14p^TPA^ variants and seamless GFP-Mdh3p. Scale bar, 5 µm. (**C**) Quantification of the cytosolic fluorescence in seamless GFP-Mdh3p cells expressing Pex14p^TPA^-wild-type, -S266A and -S266D grown in oleic acid as exemplarily shown in (B). p-values were calculated using the Welch’s test (n = 33 for WT, n = 48 for S266A, n = 20 for S266D).

**Supplementary Table S1.**
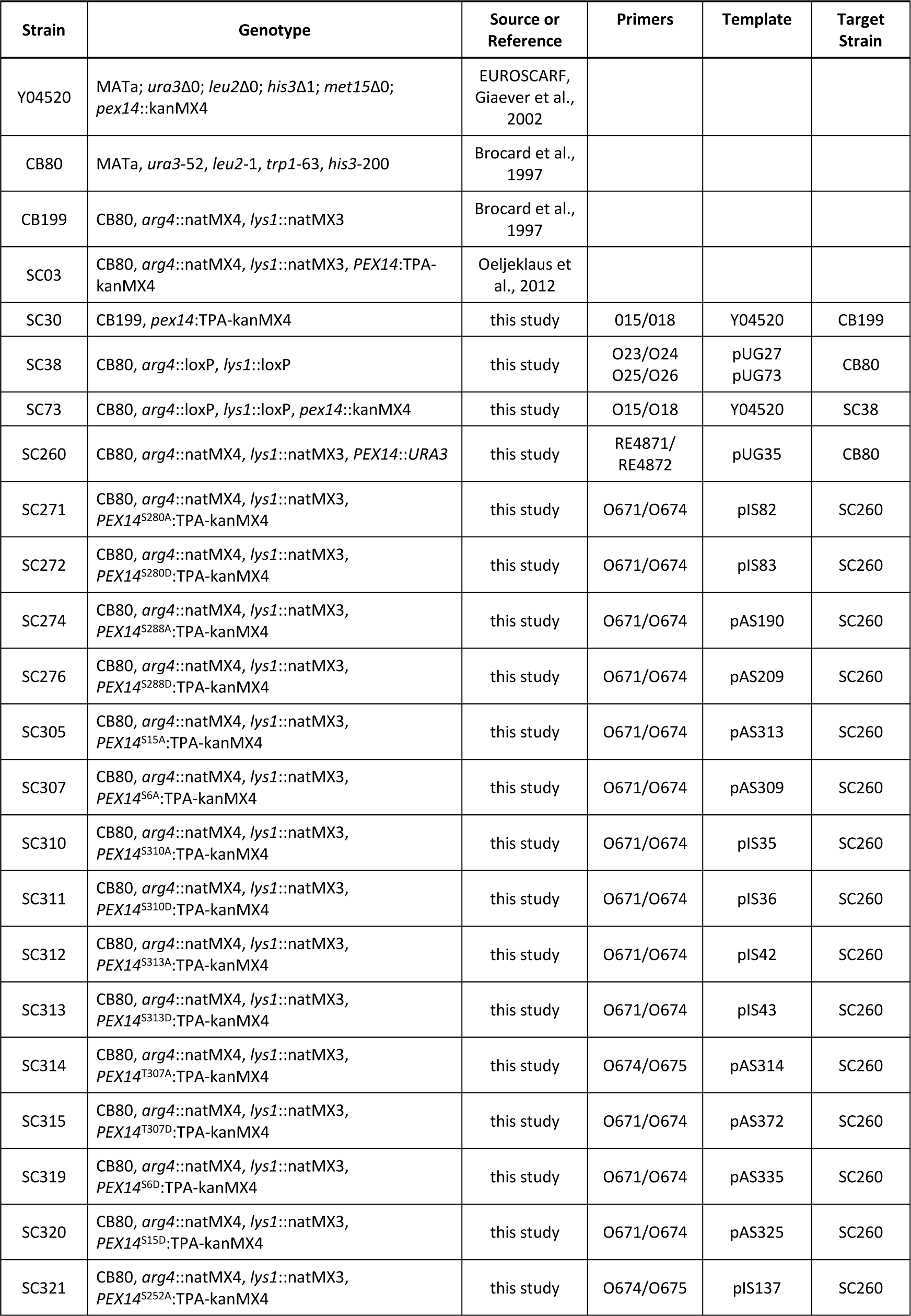
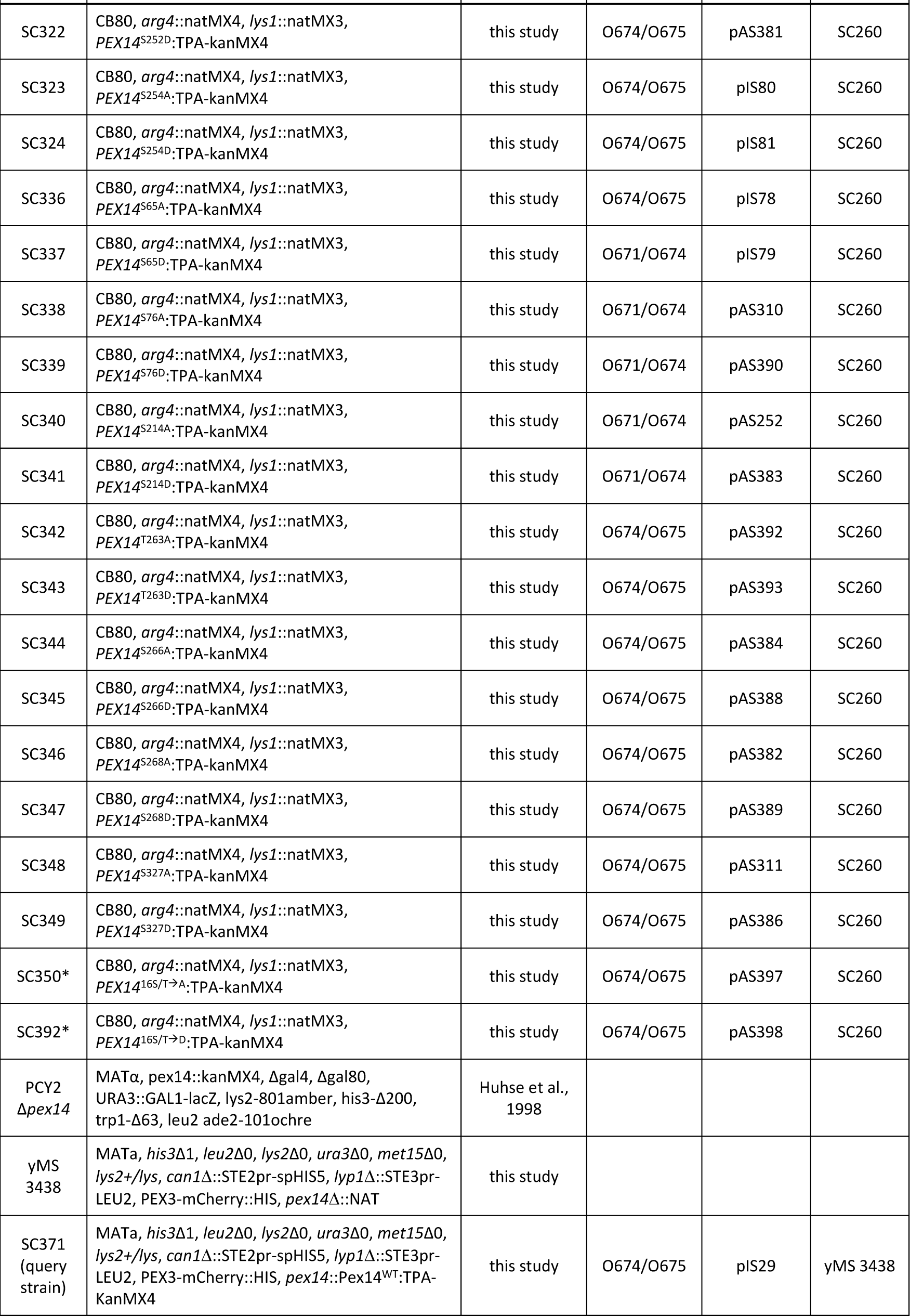
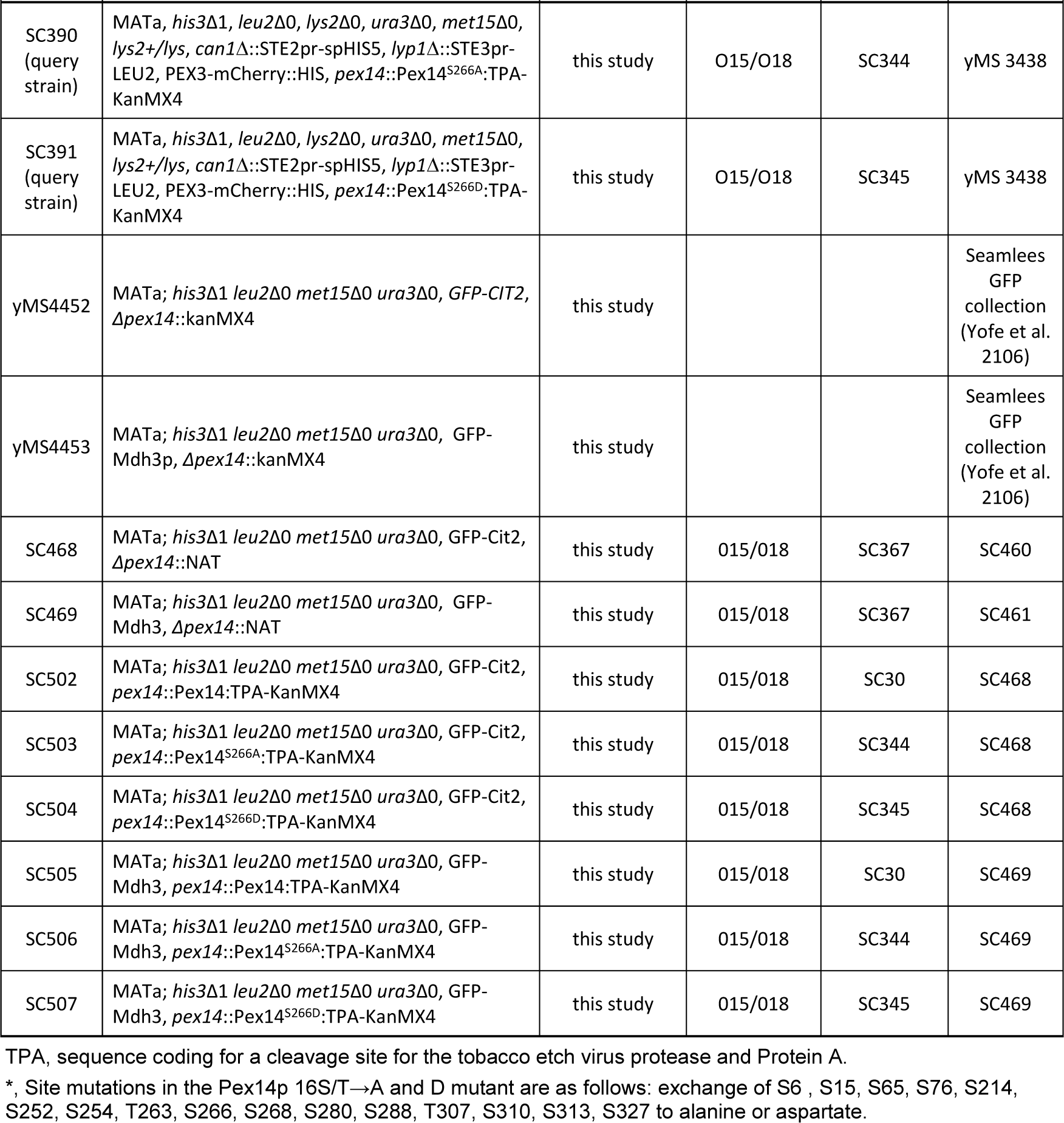
Yeast strains used in this study.

**Supplementary Table S2.**
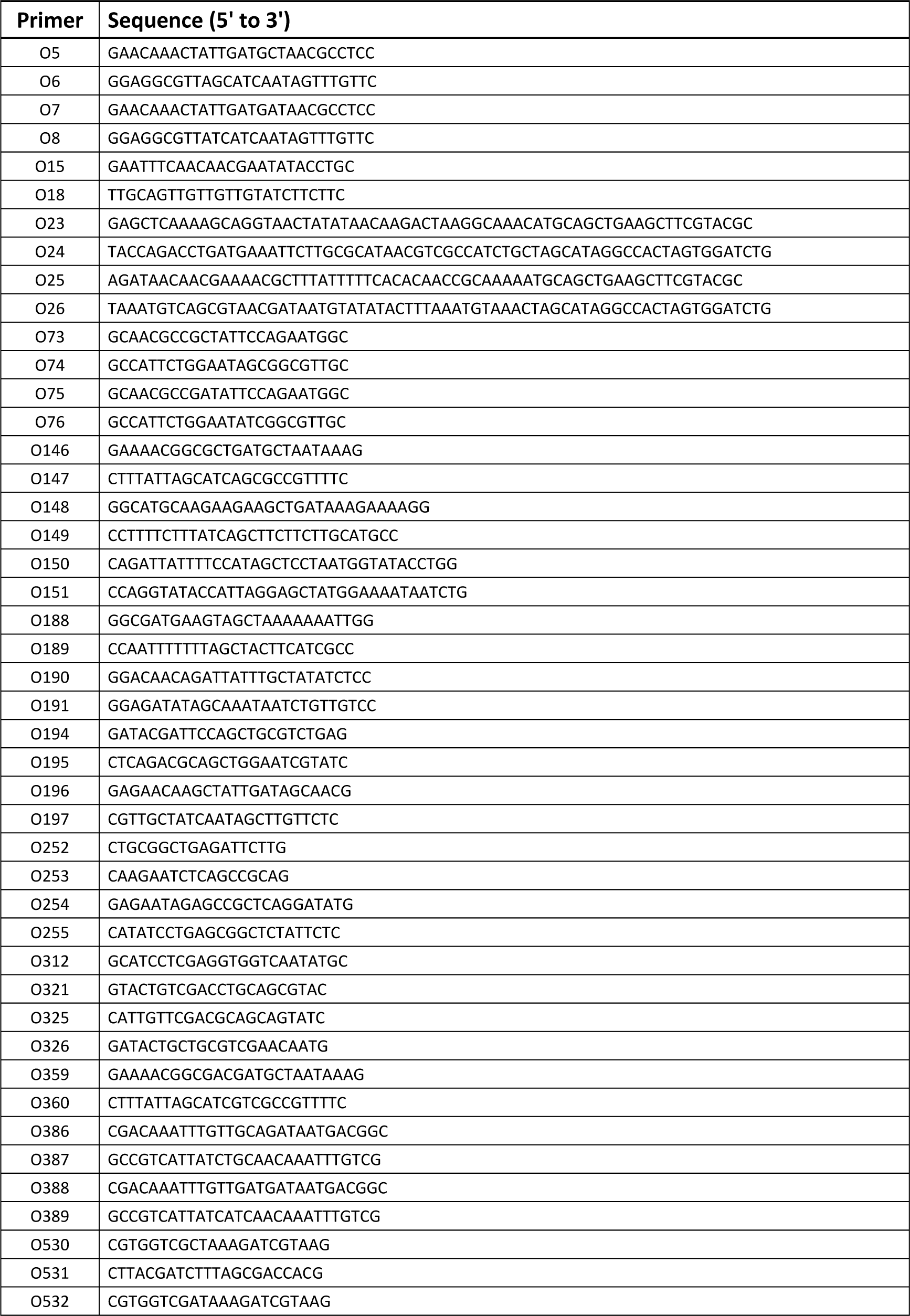
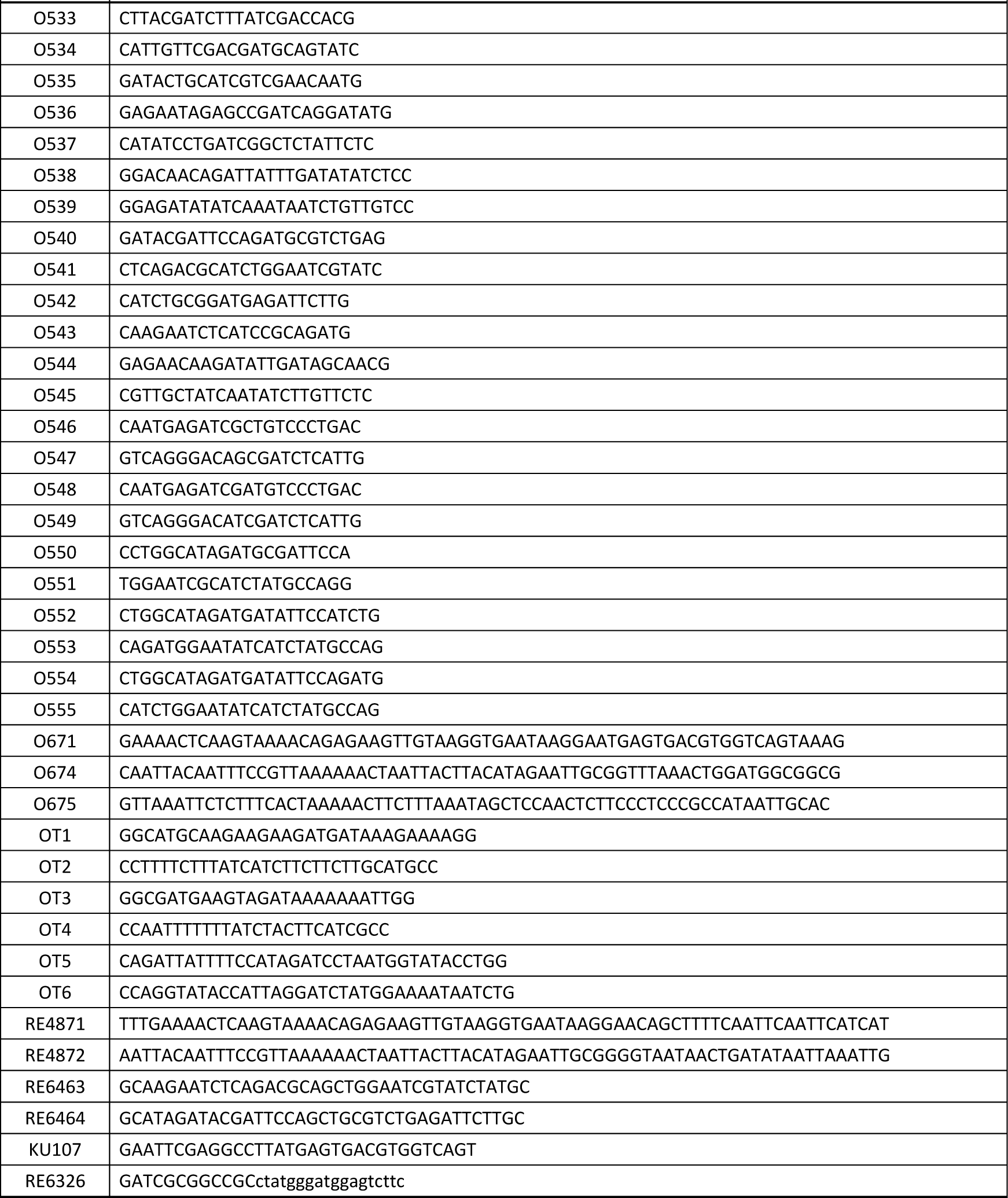
Primers used in this study.

**Supplementary Table S3.** Plasmids and cloning strategies used in this study.

| Plasmid | Characteristics/Insert/<br>Comment | Backbone | Source or<br>Reference | Primers | Restriction<br>Enzyme | Template | Target<br>Plasmid |
| --- | --- | --- | --- | --- | --- | --- | --- |
| pRS316 | *CEN6, ARSH4, URA3, bla, lacZ* | pBlue-script KS | Sikorski and Hieter, 1989 |  |  |  |  |
| pRS416 | *CEN6, ARSH4, URA3, bla, lacZ* | pBlue-script II | Sikorski and Hieter, 1989 |  |  |  |  |
| pUG27 | *loxP-AgTEF1<sub>Pro</sub>-SpHIS5-AgTEF1<sub>Term</sub>-loxP, bla* | pUG6 | Güldener et al., 2002 |  |  |  |  |
| pUG35 | *CEN6, ARSH4, URA3, MET25<sub>Pro</sub>-EGFP-Cyc1<sub>Term</sub>* | pRS416 | Niedenthal et al., 1996 |  |  |  |  |
| pUG73 | *loxP-KILEU2<sub>Pro</sub>-KILEU2-KILEU2<sub>Term</sub>-loxP, bla* | pUG6 | Güldener et al., 2002 |  |  |  |  |
| pSH47 | *CEN6, ARSH4, URA3, GAL1<sub>Pro</sub>-Cre-CYC<sub>Term</sub>, bla* | p416 | Güldener et al., 1996 |  |  |  |  |
| pGFP-SKL | *Met25<sub>Pro</sub>-GFP-SKL-Cyc1<sub>Term</sub>* | pRS416 | Schäfer et al., 2004 |  |  |  |  |
| pRS316-PEX14 | *PEX14<sub>Pro</sub>-PEX14-PEX14<sub>Term</sub>* | pRS316 | Albertini et al., 1997 |  |  |  |  |
| pIS24 | *PEX14<sub>Pro</sub>-PEX14-TPA* | pRS316 | this study | O15/O18 | EcoRI/ XhoI | gDNA* | pRS316-PEX14 |
| pIS29 | *PEX14<sub>Pro</sub>-PEX14-TPA-ADH1<sub>Term</sub>-KanMX* | pRS316 | this study | O15/O18 | EcoRI | gDNA* | pIS24 |
| pIS35 | *PEX14<sub>Pro</sub>-PEX14<sup>S310A</sup>-TPA-ADH1<sub>Term</sub>-KanMX* | pRS316 | this study | O5/O6<sup>a</sup> |  | pIS29 | pIS29 |
| pIS36 | *PEX14<sub>Pro</sub>-PEX14<sup>S310D</sup>-TPA-ADH1<sub>Term</sub>-KanMX* | pRS316 | this study | O7/O8<sup>a</sup> |  | pIS29 | pIS29 |
| pIS42 | *PEX14<sub>Pro</sub>-PEX14<sup>S313A</sup>-TPA-ADH1<sub>Term</sub>-KanMX* | pRS316 | this study | O73/O74<sup>a</sup> |  | pIS29 | pIS29 |
| pIS43 | *PEX14<sub>Pro</sub>-PEX14<sup>S313D</sup>-TPA-ADH1<sub>Term</sub>-KanMX* | pRS316 | this study | O75/O76<sup>a</sup> |  | pIS29 | pIS29 |
| pIS78 | *PEX14<sub>Pro</sub>-PEX14<sup>S65A</sup>-TPA-ADH1<sub>Term</sub>-KanMX* | pRS316 | this study | O188/O189<sup>a</sup> |  | pIS29 | pIS29 |
| pIS79 | *PEX14<sub>Pro</sub>-PEX14<sup>S65D</sup>-TPA-ADH1<sub>Term</sub>-KanMX* | pRS316 | this study | OT3/OT4<sup>a</sup> |  | pIS29 | pIS29 |
| pIS80 | *PEX14<sub>Pro</sub>-PEX14<sup>S254A</sup>-TPA-ADH1<sub>Term</sub>-KanMX* | pRS316 | this study | O150/O151<sup>a</sup> |  | pIS29 | pIS29 |
| pIS81 | *PEX14<sub>Pro</sub>-PEX14<sup>S254D</sup>-TPA-ADH1<sub>Term</sub>-KanMX* | pRS316 | this study | OT5/OT6<sup>a</sup> |  | pIS29 | pIS29 |
| pIS82 | *PEX14<sub>Pro</sub>-PEX14<sup>S280A</sup>-TPA-ADH1<sub>Term</sub>-KanMX* | pRS316 | this study | O148/O149<sup>a</sup> |  | pIS29 | pIS29 |
| pIS83 | *PEX14<sub>Pro</sub>-PEX14<sup>S280D</sup>-TPA-ADH1<sub>Term</sub>-KanMX* | pRS316 | this study | OT1/ OT2<sup>a</sup> |  | pIS29 | pIS29 |
| pIS137 | *PEX14<sub>Pro</sub>-PEX14<sup>S252A</sup>-TPA-ADH1<sub>Term</sub>-KanMX* | pRS316 | this study | O190/O191<sup>a</sup> |  | pIS29 | pIS29 |
| pAS190 | *PEX14<sub>Pro</sub>-PEX14<sup>S288A</sup>-TPA-ADH1<sub>Term</sub>-KanMX* | pRS316 | this study | O146/O147<sup>a</sup> |  | pIS29 | pIS29 |
| pAS209 | *PEX14<sub>Pro</sub>-PEX14<sup>S288D</sup>-TPA-ADH1<sub>Term</sub>-KanMX* | pRS316 | this study | O359/O360<sup>a</sup> |  | pIS29 | pIS29 |
| pAS252 | *PEX14<sub>Pro</sub>-PEX14<sup>S214A</sup>-TPA-ADH1<sub>Term</sub>-KanMX* | pRS316 | this study | O386/O387<sup>a</sup> |  | pIS29 | pIS29 |
| pAS309 | *PEX14<sub>Pro</sub>-PEX14<sup>S6A</sup>-TPA-ADH1<sub>Term</sub>-KanMX* | pRS316 | this study | O530/O531<sup>a</sup> |  | pIS29 | pIS29 |
| pAS310 | *PEX14<sub>Pro</sub>-PEX14<sup>S76A</sup>-TPA-ADH1<sub>Term</sub>-KanMX* | pRS316 | this study | O254/O255<sup>a</sup> |  | pIS29 | pIS29 |
| pAS311 | *PEX14<sub>Pro</sub>-PEX14<sup>S327A</sup>-TPA-ADH1<sub>Term</sub>-KanMX* | pRS316 | this study | O546/O547<sup>a</sup> |  | pIS29 | pIS29 |
| pAS313 | *PEX14<sub>Pro</sub>-PEX14<sup>S15A</sup>-TPA-ADH1<sub>Term</sub>-KanMX* | pRS316 | this study | O325/O326<sup>a</sup> |  | pIS29 | pIS29 |
| pAS314 | *PEX14<sub>Pro</sub>-PEX14<sup>T307A</sup>-TPA-ADH1<sub>Term</sub>-KanMX* | pRS316 | this study | O196/O197<sup>a</sup> |  | pIS29 | pIS29 |
| pAS325 | *PEX14<sub>Pro</sub>-PEX14<sup>S15D</sup>-TPA-ADH1<sub>Term</sub>-KanMX* | pRS316 | this study | O534/O535<sup>a</sup> |  | pIS29 | pIS29 |
| pAS335 | *PEX14<sub>Pro</sub>-PEX14<sup>S6D</sup>-TPA-ADH1<sub>Term</sub>-KanMX* | pRS316 | this study | O532/O533<sup>a</sup> |  | pIS29 | pIS29 |
| pAS372 | *PEX14<sub>Pro</sub>-PEX14<sup>T307D</sup>-TPA-ADH1<sub>Term</sub>-KanMX* | pRS316 | this study | O544/O545<sup>a</sup> |  | pIS29 | pIS29 |
| pAS381 | *PEX14<sub>Pro</sub>-PEX14<sup>S252D</sup>-TPA-ADH1<sub>Term</sub>-KanMX* | pRS316 | this study | O538/O539<sup>a</sup> |  | pIS29 | pIS29 |
| pAS382 | *PEX14<sub>Pro</sub>-PEX14<sup>S268A</sup>-TPA-ADH1<sub>Term</sub>-KanMX* | pRS316 | this study | O252/O253<sup>a</sup> |  | pIS29 | pIS29 |
| pAS383 | *PEX14<sub>Pro</sub>-PEX14<sup>S214D</sup>-TPA-ADH1<sub>Term</sub>-KanMX* | pRS316 | this study | O388/O389<sup>a</sup> |  | pIS29 | pIS29 |
| pAS384 | *PEX14<sub>Pro</sub>-PEX14<sup>S266A</sup>-TPA-ADH1<sub>Term</sub>-KanMX* | pRS316 | this study | O194/O195<sup>a</sup> |  | pIS29 | pIS29 |
| pAS386 | *PEX14<sub>Pro</sub>-PEX14<sup>S327D</sup>-TPA-ADH1<sub>Term</sub>-KanMX* | pRS316 | this study | O548/O549<sup>a</sup> |  | pIS29 | pIS29 |
| pAS388 | *PEX14<sub>Pro</sub>-PEX14<sup>S266D</sup>-TPA-ADH1<sub>Term</sub>-KanMX* | pRS316 | this study | O540/O541<sup>a</sup> |  | pIS29 | pIS29 |
| pAS389 | *PEX14<sub>Pro</sub>-PEX14<sup>S268D</sup>-TPA-ADH1<sub>Term</sub>-KanMX* | pRS316 | this study | O542/O543<sup>a</sup> |  | pIS29 | pIS29 |
| pAS390 | *PEX14<sub>Pro</sub>-PEX14<sup>S76D</sup>-TPA-ADH1<sub>Term</sub>-KanMX* | pRS316 | this study | O536/O537<sup>a</sup> |  | pIS29 | pIS29 |
| pAS392 | *PEX14<sub>Pro</sub>-PEX14<sup>T263A</sup>-TPA-ADH1<sub>Term</sub>-KanMX* | pRS316 | this study | O550/O551<sup>a</sup> |  | pIS29 | pIS29 |
| pAS393 | *PEX14<sub>Pro</sub>-PEX14<sup>T263D</sup>-TPA-ADH1<sub>Term</sub>-KanMX* | pRS316 | this study | O552/O553<sup>a</sup> |  | pIS29 | pIS29 |
| pAS335 | *PEX14<sub>Pro</sub>-PEX14<sup>S6D</sup>-TPA-ADH1<sub>Term</sub>-KanMX* | pRS316 | this study | O532/O533<sup>a</sup> |  | pIS29 | pIS29 |
| pAS372 | *PEX14<sub>Pro</sub>-PEX14<sup>T307D</sup>-TPA-ADH1<sub>Term</sub>-KanMX* | pRS316 | this study | O544/O545<sup>a</sup> |  | pIS29 | pIS29 |
| pAS381 | *PEX14<sub>Pro</sub>-PEX14<sup>S252D</sup>-TPA-ADH1<sub>Term</sub>-KanMX* | pRS316 | this study | O538/O539<sup>a</sup> |  | pIS29 | pIS29 |
| pAS382 | *PEX14<sub>Pro</sub>-PEX14<sup>S268A</sup>-TPA-ADH1<sub>Term</sub>-KanMX* | pRS316 | this study | O252/O253<sup>a</sup> |  | pIS29 | pIS29 |
| pAS383 | *PEX14<sub>Pro</sub>-PEX14<sup>S214D</sup>-TPA-ADH1<sub>Term</sub>-KanMX* | pRS316 | this study | O388/O389<sup>a</sup> |  | pIS29 | pIS29 |
| pAS384 | *PEX14<sub>Pro</sub>-PEX14<sup>S266A</sup>-TPA-ADH1<sub>Term</sub>-KanMX* | pRS316 | this study | O194/O195<sup>a</sup> |  | pIS29 | pIS29 |
| pAS386 | *PEX14<sub>Pro</sub>-PEX14<sup>S327D</sup>-TPA-ADH1<sub>Term</sub>-KanMX* | pRS316 | this study | O548/O549<sup>a</sup> |  | pIS29 | pIS29 |
| pAS388 | *PEX14<sub>Pro</sub>-PEX14<sup>S266D</sup>-TPA-ADH1<sub>Term</sub>-KanMX* | pRS316 | this study | O540/O541<sup>a</sup> |  | pIS29 | pIS29 |
| pAS389 | *PEX14<sub>Pro</sub>-PEX14<sup>S268D</sup>-TPA-ADH1<sub>Term</sub>-KanMX* | pRS316 | this study | O542/O543<sup>a</sup> |  | pIS29 | pIS29 |
| pAS390 | *PEX14<sub>Pro</sub>-PEX14<sup>S76D</sup>-TPA-ADH1<sub>Term</sub>-KanMX* | pRS316 | this study | O536/O537<sup>a</sup> |  | pIS29 | pIS29 |
| pAS392 | *PEX14<sub>Pro</sub>-PEX14<sup>T263A</sup>-TPA-ADH1<sub>Term</sub>-KanMX* | pRS316 | this study | O550/O551<sup>a</sup> |  | pIS29 | pIS29 |
| pAS393 | *PEX14<sub>Pro</sub>-PEX14<sup>T263D</sup>-TPA-ADH1<sub>Term</sub>-KanMX* | pRS316 | this study | O552/O553<sup>a</sup> |  | pIS29 | pIS29 |
| pAS397<sup>b,c</sup> | *PEX14<sub>Pro</sub>-PEX14<sup>16A</sup>-TPA-ADH1<sub>Term</sub>-KanMX* | pRS316 | this study | O188/O189<sup>a</sup><br>O254/O255<sup>a</sup><br>O325/O326<sup>a</sup><br>O386/O387<sup>a</sup><br>O530/O531<sup>a</sup><br>O546/O547<sup>a</sup><br>O550/O551<sup>a</sup><br>O312/O321 | XhoI/SalI | pIS29,<br>GeneArt<br>string<sup>d</sup> | pIS29 |
| pAS398<sup>b,e</sup> | *PEX14<sub>Pro</sub>-PEX14<sup>16D</sup>-TPA-ADH1<sub>Term</sub>-KanMX* | pRS316 | this study | O388/O389<sup>a</sup><br>O532/O533<sup>a</sup><br>O548/O549<sup>a</sup><br>O554/O555<sup>a</sup> | XbaI/SalI | pEX-K-PEX14<sup>f</sup> | pIS29 |
| pEX-K-PEX14<sup>f</sup> | *Kan, pUC ori, PEX14<sub>Pro</sub>-PEX14<sup>12D</sup>* |  | Eurofins,<br>this study |  |  | Gene<br>synthesis |  |
| pPC86 | *ADC1<sub>Pro</sub>-Gal4-activation*<br>*domain (Gal4-AD)* |  | Chevray<br>and<br>Nathans,<br>1992 |  |  |  |  |
| pPC97 | *ADC1<sub>Pro</sub>-Gal4-DNA*<br>*binding domain (Gal4-BD)* |  | Chevray<br>and<br>Nathans,<br>1992 |  |  |  |  |
| pPC86 +<br>PEX5 | *ADC1<sub>Pro</sub>-Gal4-AD-PEX5* | pPC86 | Erdmann<br>and Blobel,<br>1996 |  |  |  |  |
| pPC97 +<br>PEX13 | *ADC1<sub>Pro</sub>-Gal4-BD-PEX13* | pPC97 | Girzalsky<br>et al., 1999 |  |  |  |  |
| pPC86 +<br>PEX14 | *ADC1<sub>Pro</sub>-Gal4-AD-PEX14* | pPC86 | Albertini et<br>al., 1997 |  |  |  |  |
| pPC97 +<br>PEX14 | *ADC1<sub>Pro</sub>-Gal4-BD-PEX14* | pPC97 | Albertini et<br>al., 1997 |  |  |  |  |
| pPC97 +<br>PEX17 | *ADC1<sub>PRO</sub>-Gal4-BD-PEX17* | pPC97 | Albertini et<br>al., 1997 |  |  |  |  |
| pPC86 +<br>PEX14<sup>S266D</sup> | *ADC1<sub>PRO</sub>-Gal4-AD-*<br>*PEX14<sup>S266D</sup>* | pPC86 | this study | KU107/<br>RE6326 | EcoRI/NotI | gDNA |  |
| pPC86 +<br>PEX14<sup>S266A</sup> | *ADC1<sub>PRO</sub>-Gal4-AD-*<br>*PEX14<sup>S266A</sup>* | pPC86 | this study | RE6463/<br>RE6464 |  | pPC86 +<br>PEX14 |  |
| pPC97 +<br>PEX14<sup>S266D</sup> | *ADC1<sub>PRO</sub>-Gal4-BD-*<br>*PEX14<sup>S266D</sup>* | pPC97 | this study |  | Sall/NotI | pPC86 +<br>PEX14<sup>S266D</sup> |  |
| pPC97 +<br>PEX14<sup>S266A</sup> | *ADC1<sub>PRO</sub>-Gal4-BD-*<br>*PEX14<sup>S266A</sup>* | pPC97 | this study | RE6463/<br>RE6464 |  | pPC97 +<br>PEX14 |  |

Pro, promoter; Term, terminator; TPA, sequence coding for a cleavage site for the tobacco etch virus protease and Protein A
\* Genomic DNA (gDNA) was prepared from SC03
<sup>a</sup>, Used for site-directed mutagenesis according to Papworth et al. (1996).
<sup>b</sup>, Site mutations in the Pex14p-16A and -16D mutants are as follows: exchange of S6, S15, S65, S76, S214, S252, S254, T263, S266, S268, S280, S288, T307, S310, S313, S327 to alanine (16A) or aspartate (16D).
<sup>c</sup>, For generation of the Pex14p-16A mutant, S6 (O530/O531), S15 (O325/O326), S65 (O188/O189), S76 (O254/O255), S214 (O386/O387), T263 (O550/O551), and S327 (O546/O547) to Ala mutations were introduced by consecutive site-directed mutageneses. The remaining 9 site mutations were introduced using gene synthesis (GeneArt Strings DNA Fragments, Thermo Fisher Scientific), amplified by PCR (O312/O321) and inserted in XhoI/Sall sites.
<sup>e</sup>, For generation of the Pex14p-16D mutant, S6 (O532/O533), S214 (O388/O389), T263 (O554/O555), and S327 (O548/O549) to Asp mutations were introduced by consecutive site-directed mutageneses. The remaining 12 site mutations were introduced using gene synthesis (Eurofins), integrated in the vector pEX-K and ligated in the target vector in XbaI/Sall sites.

## Notes

### Competing Interest Statement

The authors have declared no competing interest.

