## Supplementary material for "Pex14p phosphorylation modulates import of citrate synthase 2 into peroxisomes in *Saccharomyces cerevisiae*": Figure S2

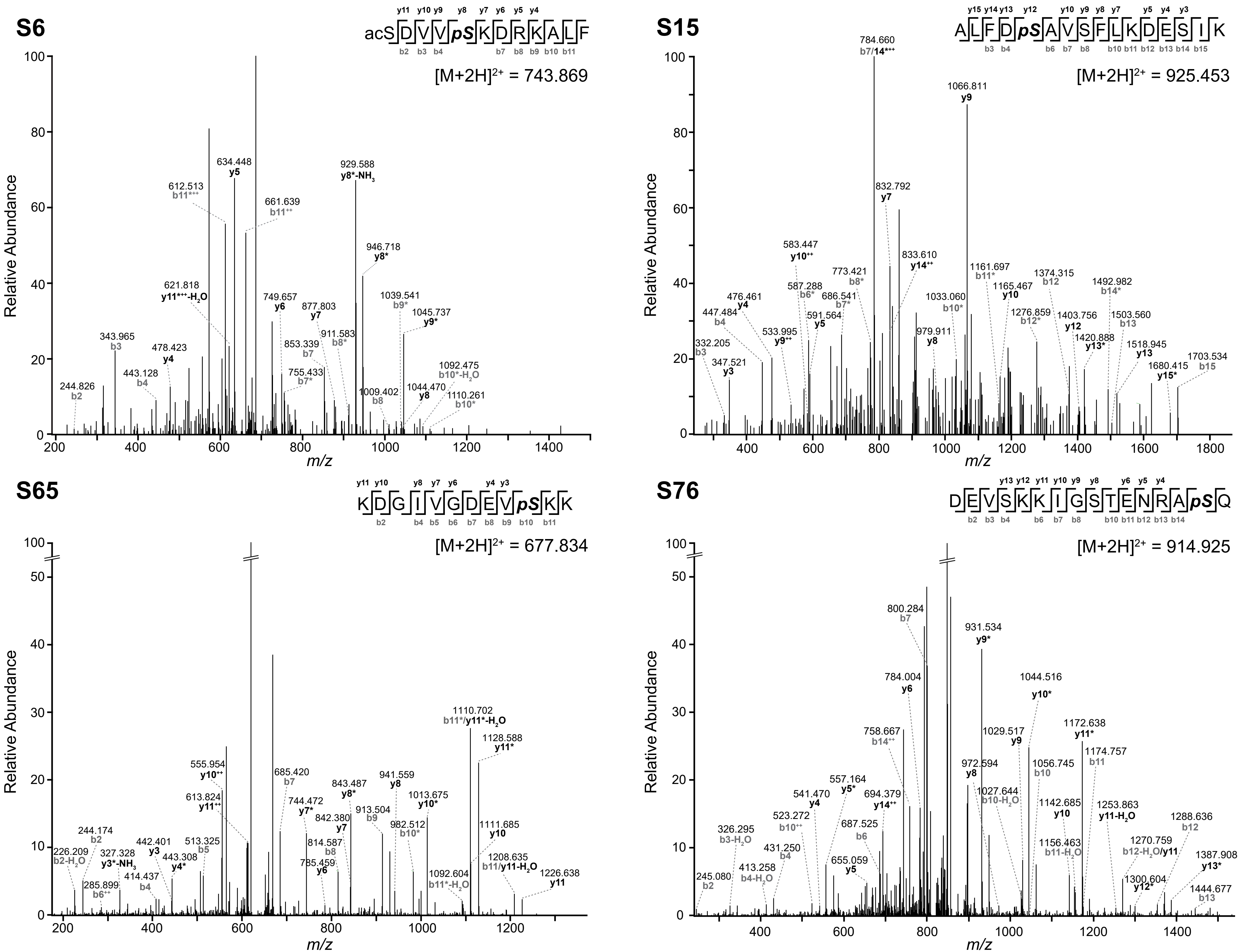
S6
acSDVVpSKDRKALF
[M+2H]2+ = 743.869
Relative Abundance
m/z
634.448 y5
929.588 y8*-NH3
612.513 b11+++
661.639 b11++
946.718 y8*
621.818 y11+++-H2O
749.657 y6
1039.541 b9*
877.803 y7
1045.737 y9*
343.965 b3
478.423 y4
853.339 b7
911.583 b8*
443.128 b4
1092.475 b10*-H2O
755.433 b7*
244.826 b2
1044.470 y8
1110.261 b10*
1009.402 b8
S15
ALFDpSAVSFLKDESIK
[M+2H]2+ = 925.453
784.660 b7/14+++
1066.811 y9
832.792 y7
583.447 y10++
833.610 y14++
773.421 b8*
1161.697 b11*
1374.315 b12
587.288 b6*
1165.467 y10
1492.982 b14*
476.461 y4
686.541 b7*
1033.060 b10*
447.484 b4
1276.859 b12*
1403.756 y12
1503.560 b13
533.995 y9++
591.564 y5
979.911 y8
1420.888 y13*
332.205 b3
1518.945 y13
347.521 y3
1703.534 b15
1680.415 y15*
S65
KDGIVGDEVpSKK
[M+2H]2+ = 677.834
1110.702 b11*/y11*-H2O
1128.588 y11*
555.954 y10++
685.420 b7
941.559 y8
613.824 y11++
843.487 y8*
1013.675 y10*
744.472 y7*
913.504 b9
842.380 y7
982.512 b10*
1111.685 y10
244.174 b2
814.587 b8
442.401 y3
513.325 b5
1208.635 b11/y11-H2O
226.209 b2-H2O
327.328 y3*-NH3
443.308 y4*
785.459 y6
1226.638 y11
414.437 b4
285.899 b6++
1092.604 b11*-H2O
S76
DEVSKKIGSTENRApSQ
[M+2H]2+ = 914.925
800.284 b7
931.534 y9*
784.004 y6
1044.516 y10*
1172.638 y11*
758.667 b14++
1029.517 y9
1174.757 b11
972.594 y8
1056.745 b10
557.164 y5*
541.470 y4
694.379 y14++
1027.644 b10-H2O
1142.685 y10
1253.863 y11-H2O
1288.636 b12
523.272 b10++
687.525 b6
326.295 b3-H2O
431.250 b4
1270.759 b12-H2O/y11
1156.463 b11-H2O
1387.908 y13*
655.059 y5
413.258 b4-H2O
245.080 b2
1300.604 y12*
1444.677 b13

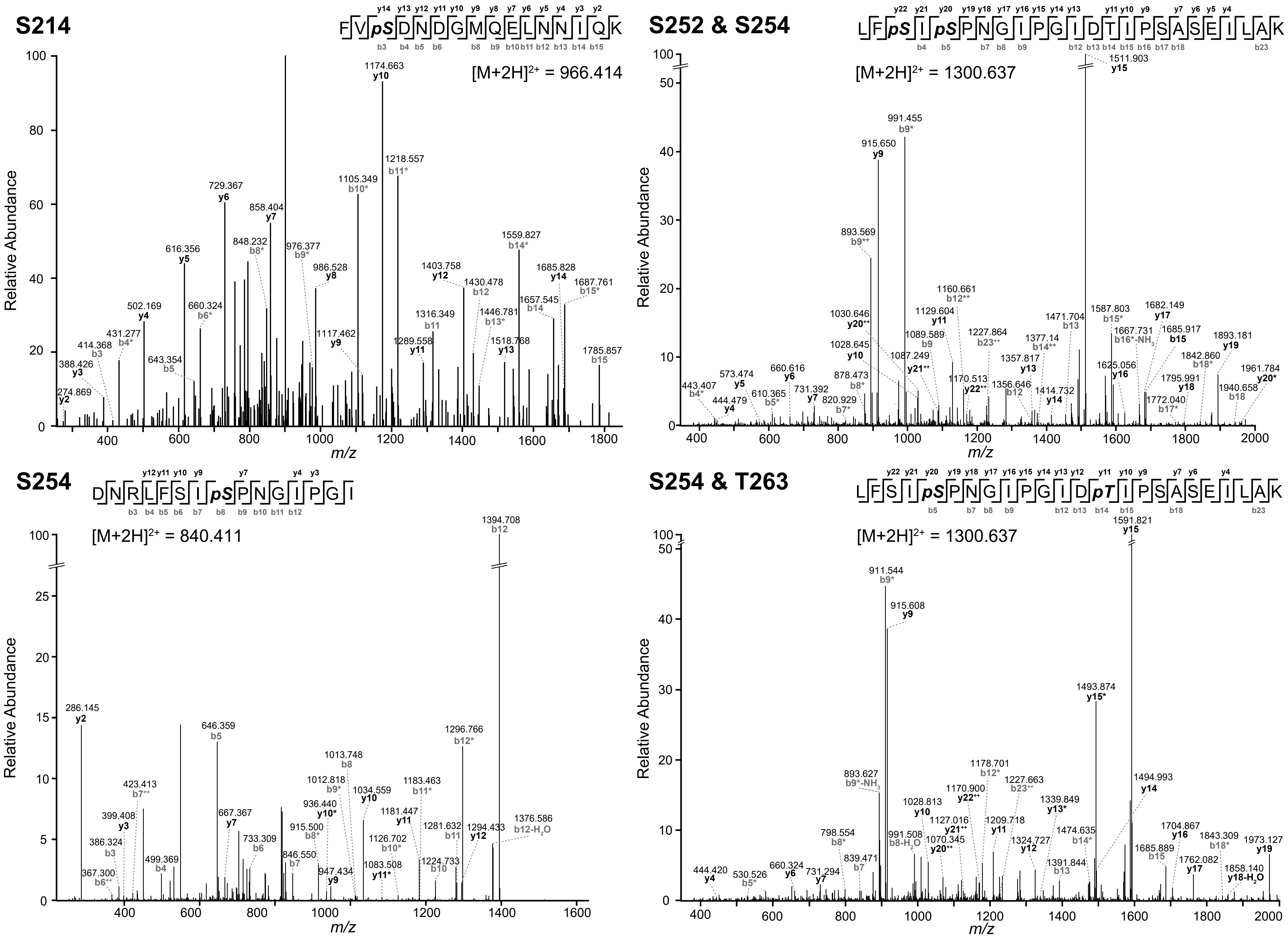
S214
FVpSDNDGMQELNNIQK
[M+2H]$^{2+}$ = 966.414
Relative Abundance
m/z
1174.663 y10
1218.557 b11*
1105.349 b10*
729.367 y6
858.404 y7
848.232 b8*
976.377 b9*
986.528 y8
616.356 y5
1403.758 y12
1430.478 b12
1559.827 b14*
1685.828 y14
1687.761 b15*
1657.545 b14
1446.781 b13*
1316.349 b11
502.169 y4
660.324 b6*
431.277 b4*
414.368 b3
388.426 y3
274.869 y2
643.354 b5
1117.462 y9
1289.558 y11
1518.768 y13
1785.857 b15
S252 & S254
LFpSIpSPNGIPGIDTIPSASEILAK
[M+2H]$^{2+}$ = 1300.637
1511.903 y15
991.455 b9*
915.650 y9
893.569 b9**
1160.661 b12**
1129.604 y11
1030.646 y20$^{++}$
1089.589 b9
1227.864 b23$^{++}$
1471.704 b13
1587.803 b15*
1682.149 y17
1667.731 b16*-NH$_3$
1685.917 b15
1893.181 y19
1028.645 y10
1087.249 y21$^{++}$
1357.817 y13
1377.14 b14**
1625.056 y16
1842.860 b18*
573.474 y5
660.616 y6
878.473 b8*
1170.513 y22$^{++}$
1356.646 b12
1414.732 y14
1795.991 y18
1940.658 b18
1961.784 y20*
443.407 b4*
444.479 y4
610.365 b5*
731.392 y7
820.929 b7*
1772.040 b17*
S254
DNRLFSIpSPNGIPGI
[M+2H]$^{2+}$ = 840.411
1394.708 b12
286.145 y2
646.359 b5
1296.766 b12*
1013.748 b8
1012.818 b9*
1034.559 y10
1183.463 b11*
423.413 b7$^{++}$
936.440 y10*
1181.447 y11
399.408 y3
667.367 y7
915.500 b8*
1281.632 b11
1294.433 y12
1376.586 b12-H$_2$O
386.324 b3
733.309 b6
1126.702 b10*
499.369 b4
846.550 b7
1083.508 y11*
1224.733 b10
367.300 b6$^{++}$
947.434 y9
S254 & T263
LFSIpSPNGIPGIDpTIPSASEILAK
[M+2H]$^{2+}$ = 1300.637
1591.821 y15
911.544 b9*
915.608 y9
1493.874 y15*
1178.701 b12*
1227.663 b23$^{++}$
893.627 b9*-NH$_3$
1170.900 y22$^{++}$
1494.993 y14
1339.849 y13*
1028.813 y10
1127.016 y21$^{++}$
1209.718 y11
1474.635 b14*
1704.867 y16
1843.309 b18*
798.554 b8*
991.508 b8-H$_2$O
1070.345 y20$^{++}$
1324.727 y12
1973.127 y19
1685.889 b15
1762.082 y17
1391.844 b13
1858.140 y18-H$_2$O
444.420 y4
530.526 b5*
660.324 y6
731.294 y7
839.471 b7

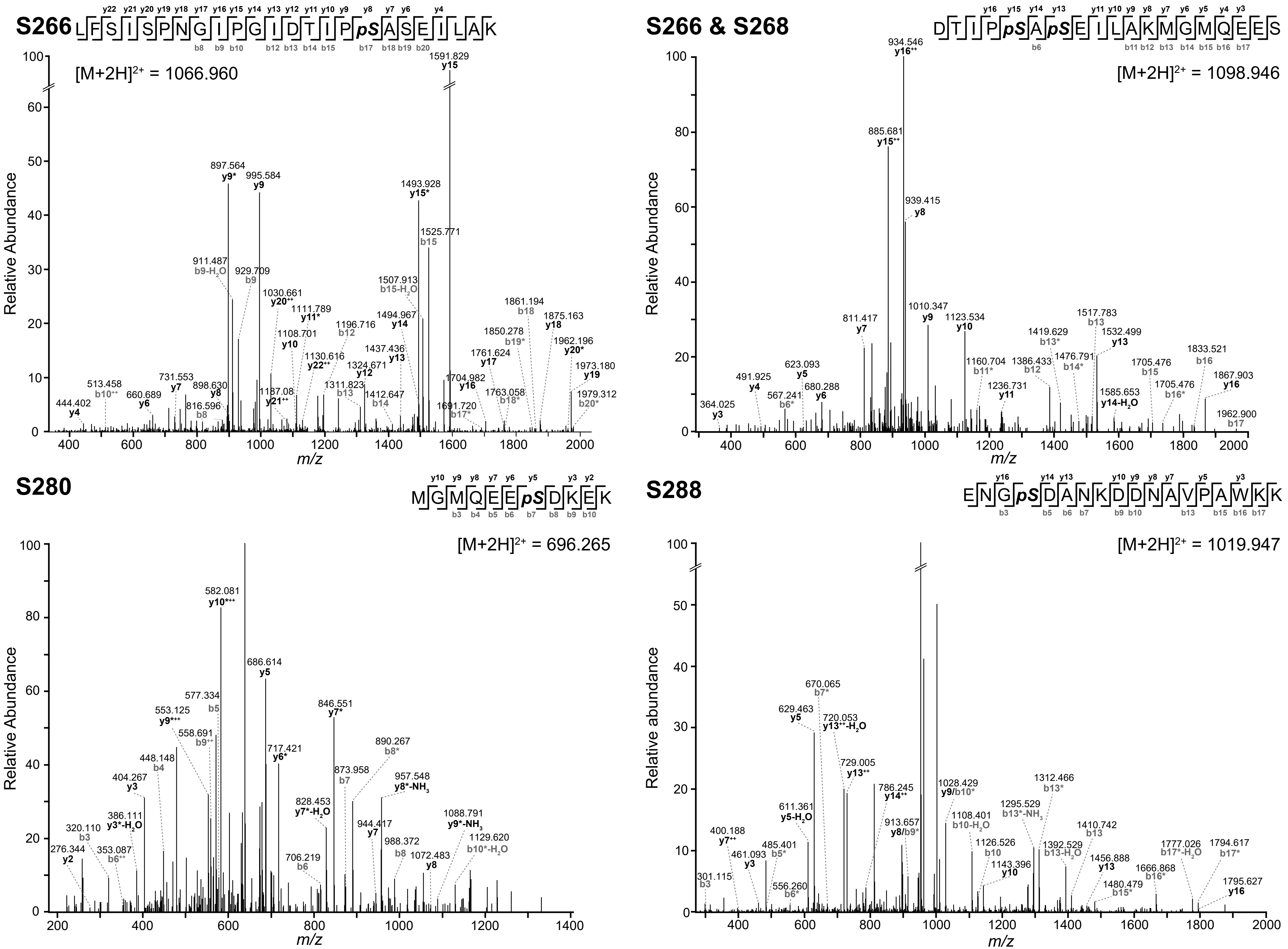
S266
LFSISPNGIPGIDTIPpSASEILAK
[M+2H]$^{2+}$ = 1066.960
1591.829 y15
897.564 y9*
995.584 y9
1493.928 y15*
1525.771 b15
911.487 b9-H$_2$O
929.709 b9
1507.913 b15-H$_2$O
1030.661 y20$^{++}$
1111.789 y11*
1494.967 y14
1861.194 b18
1875.163 y18
1196.716 b12
1108.701 y10
1850.278 b19*
1962.196 y20*
1130.616 y22$^{++}$
1437.436 y13
1761.624 y17
1324.671 y12
1973.180 y19
513.458 b10$^{++}$
731.553 y7
898.630 y8
1311.823 b13
1412.647 b14
1704.982 y16
1763.058 b18*
1979.312 b20*
660.689 y6
1187.08 y21$^{++}$
1691.720 b17*
444.402 y4
816.596 b8
S266 & S268
DTIPpSApSEILAKMGMQEES
[M+2H]$^{2+}$ = 1098.946
934.546 y16$^{++}$
885.681 y15$^{++}$
939.415 y8
1010.347 y9
1123.534 y10
811.417 y7
1517.783 b13
1532.499 y13
1419.629 b13*
1476.791 b14*
1833.521 b16
1160.704 b11*
1386.433 b12
1705.476 b15
1867.903 y16
623.093 y5
491.925 y4
680.288 y6
567.241 b6*
1236.731 y11
1585.653 y14-H$_2$O
1705.476 b16*
364.025 y3
1962.900 b17
S280
MGMQEEpSDKEK
[M+2H]$^{2+}$ = 696.265
582.081 y10$^{+++}$
686.614 y5
577.334 b5
553.125 y9$^{+++}$
558.691 b9$^{++}$
846.551 y7*
717.421 y6*
890.267 b8*
448.148 b4
404.267 y3
873.958 b7
957.548 y8*-NH$_3$
828.453 y7*-H$_2$O
386.111 y3*-H$_2$O
944.417 y7
1088.791 y9*-NH$_3$
320.110 b3
1129.620 b10*-H$_2$O
988.372 b8
276.344 y2
353.087 b6$^{++}$
706.219 b6
1072.483 y8
S288
ENGpSDANKDDNAVPAWKK
[M+2H]$^{2+}$ = 1019.947
670.065 b7*
629.463 y5
720.053 y13$^{++}$-H$_2$O
729.005 y13$^{++}$
786.245 y14$^{++}$
1028.429 y9/b10*
1312.466 b13*
1295.529 b13*-NH$_3$
611.361 y5-H$_2$O
913.657 y8/b9*
1108.401 b10-H$_2$O
1410.742 b13
400.188 y7$^{++}$
1126.526 b10
1392.529 b13-H$_2$O
1777.026 b17*-H$_2$O
1794.617 b17*
461.093 y3
485.401 b5*
1143.396 y10
1456.888 y13
1666.868 b16*
1480.479 b15*
1795.627 y16
301.115 b3
556.260 b6*
Relative Abundance
Relative abundance
m/z

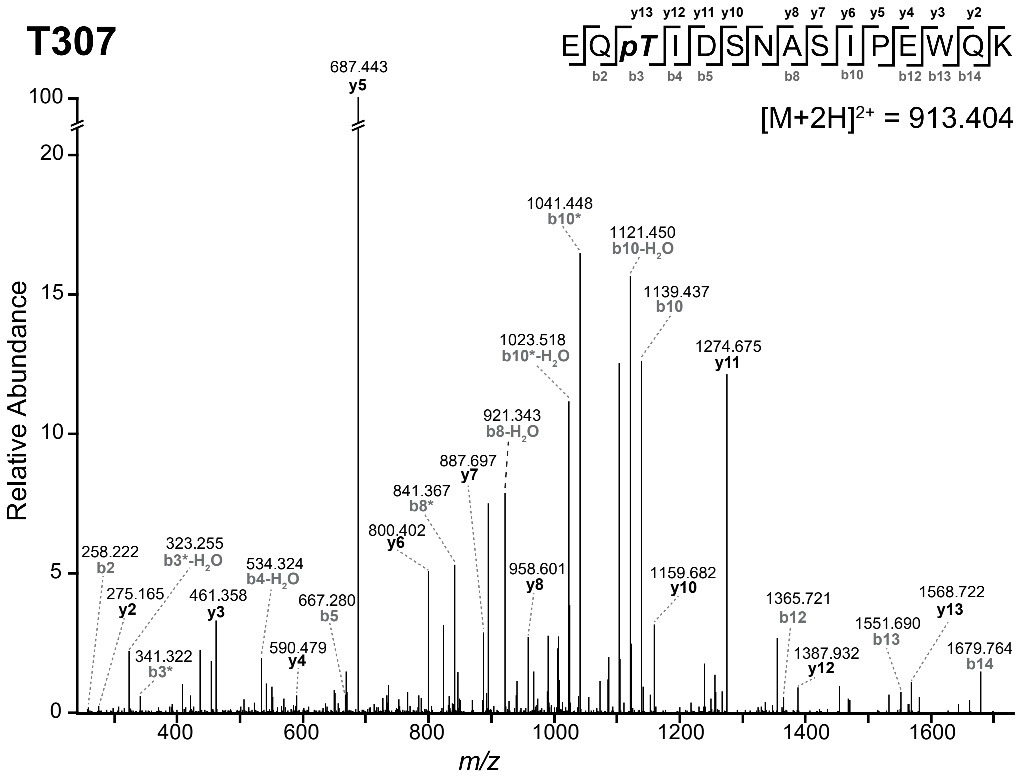


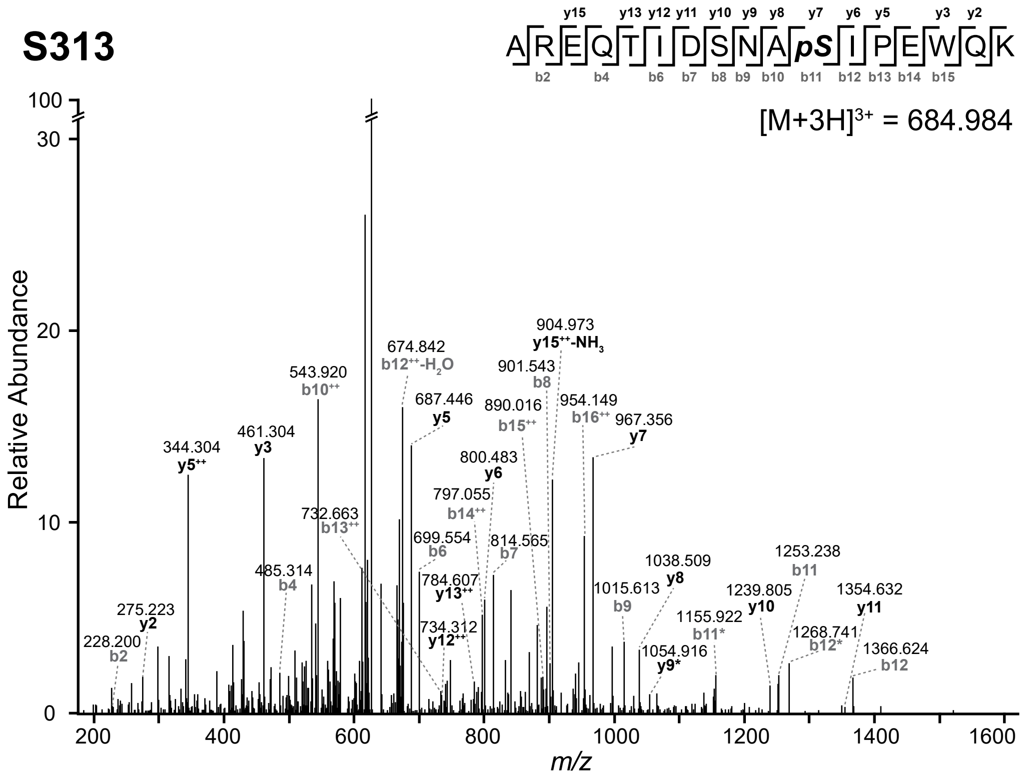


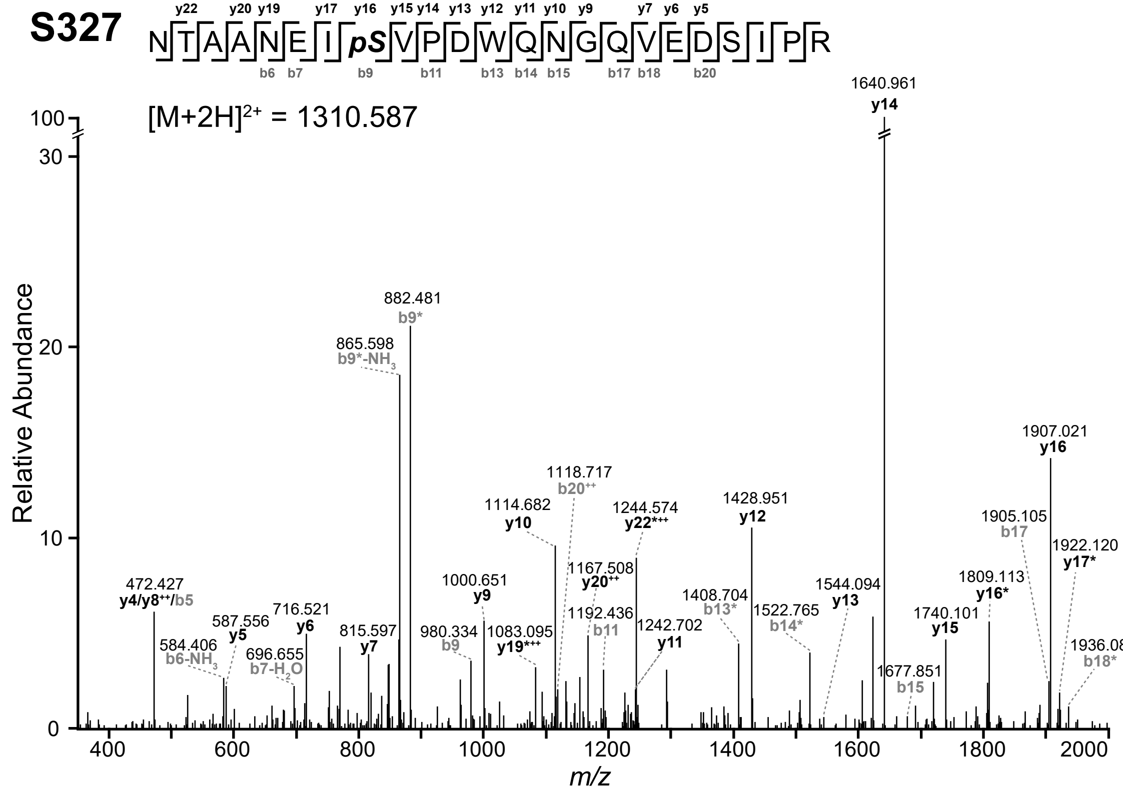


Supplementary Figure S2: Annotated MS/MS spectra of Pex14p peptide ions comprising the phosphorylation site(s) indicated in the figure. The spectra were acquired on an LTQ-Orbitrap XL using MSA for fragmentation of the precursor ion. ++, doubly protonated fragment ion; */** fragment ion with one/two neutral loss(es) of $H_3PO_4$; ac, acetylation.
